# Clinical isolates of *Enterobacter* species can persist in human macrophages without bacterial replication and minimal cellular cytotoxicity

**DOI:** 10.1101/2025.02.08.637254

**Authors:** Georgiana Parau, Hannah J. Parks, Amy J. G. Anderson, Fabiana Bisaro, Inmaculada García-Romero, Michael C. Gilmore, Samuel O. Korankye, Helina Marshall, Miguel A. Valvano

## Abstract

**Background:** The *Enterobacter cloacae* complex (Ecc) encompasses opportunistic Gram-negative bacteria demonstrating considerable phenotypic and genotypic diversity. Bloodstream, respiratory and urinary tract infections by Ecc bacteria are associated with morbidity and mortality worldwide. These infections are often difficult to treat since Ecc bacteria are resistant to penicillins, quinolones, aminoglycosides, and third-generation cephalosporins. Resistance also extends to carbapenems, leaving only polymyxins, such as colistin, as a last resort antibiotic for treatment. However, colistin resistance in Ecc isolates is also unexpectedly frequent. Despite extensive information on antibiotic resistance by *Enterobacter* species, much less is known about their infection biology. There are few reports on the survival and persistence of selected *Enterobacter* species in macrophages and epithelial cells, but how *Enterobacter* isolates interact with innate immune host cells upon engulfment remains unexplored. In this study, we have investigated the intracellular trafficking of a subset of antimicrobial resistant Ecc clinical isolates, including colistin-resistant strains, within human macrophages, and determined the macrophage response to the intracellular infection.

**Methods:** Phagocytosis of 11 clinical Ecc isolates, including *E. cloacae, E. bugandensis, E. kobei, E. xiangfangensis, E. roggenkampii, E. hoffmannii*, and *E. ludwigii* was investigated in THP-1 and human monocyte derived macrophages (HMDMs). Confocal fluorescence microscopy was used to ascertain intracellular trafficking via co-localisation of cell markers with fluorescent bacteria. Intracellular bacterial replication was assessed by bacterial enumeration in cell lysates after killing extracellular bacteria and by a fluorescence dilution approach to follow the synthesis of the bacterial cell wall over time. Macrophage cell cytotoxicity was investigated by quantifying the release of lactate dehydrogenase during infection with all isolates. Two prototypic isolates, the *E. cloacae* ATCC13047 type strain and the *E. bugandensis* 104107, were used to explore in more detail the response of macrophages to the intracellular infection by determining cleavage of the proinflammatory markers caspase-1, gasdermin D and pro-interleukin-1β.

**Findings:** We found that Ecc isolates do not replicate in human macrophages but survive within a modified late phagolysosome compartment. Survival occurred in all species investigated and did not correlate with colistin resistance, lipopolysaccharide modifications, or bacterial pathogenicity in the *Galleria mellonella* infection model. All isolates induced macrophage cell cytotoxicity at significantly lower levels than controls treated with lipopolysaccharide and nigericin treatment (to induce a proinflammatory response). Low cytotoxicity also correlated with absence of cleavage of proinflammatory markers in infected macrophages.

**Interpretation:** Ecc species can survive without replication inside human macrophages with minimal effects on cell viability and inflammation. These observations could have implications in the clinical outcome of patients that cannot readily clear infecting Ecc bacteria. This can potentially lead to prolonged intracellular survival and infection relapse.

**Funding:** Biotechnology and Biological Sciences Research Council grants BB/T005807/1 and BB/S006281/1.

**Research In context:** *Evidence before this study:* We searched PubMed with the terms “Enterobacter” AND “macrophage”, “Enterobacter” AND “monocytes”, and “Enterobacter” AND “intracellular” for original articles published in English up to July 9, 2024. The search excluded terms “sakazakii” and “aerogenes” since *E. aerogenes* and *E. sakazakii* have been moved to the genus *Klebsiella* and *Cronobacter*, respectively. Of the 55, 15, and 181 studies we identified, respectively, only one reported testing *Enterobacter cloacae* phagocytosis. Another study reported intracellular bacterial communities in uroepithelial cells, which serve as a reservoir during urinary tract infection. One other study reported the isolation of *E. hormaechei* from human atherosclerotic tissue and described studies in THP-1 monocytic macrophages. A few earlier studies also reported *Enterobacter* cytotoxins affecting immune cells, and an *E. cloacae* polysaccharide capable of inducing apoptosis in epithelial cells. These studies did not investigate mechanisms and there have been no more recent follow-ups; importantly, it remains unclear if the strains employed in these studies were properly identified as *Enterobacter* species. Therefore, despite isolated observations of survival of *Enterobacter* isolates surviving in immune and epithelial cells, there is an overall knowledge gap in our understanding of these pathogens concerning intracellular survival compartments, kinetics of survival, and induction of macrophage cytotoxicity and inflammatory responses.

*Added value of this study:* Our study is the first to investigate in detail how clinical isolates of various *Enterobacter* species can survive intracellularly in human macrophages. All isolates display multidrug antimicrobial resistance, including some with colistin resistance, and can survive intracellularly in human macrophages. Our data demonstrate that intracellular *Enterobacter* resides in vacuoles for up to 44 hours without replication. Colocalization experiments with various fluid phase and membrane cellular markers revealed that bacteria-containing vacuoles are modified late phagolysosomes, which do not accumulate the autophagosome marker LC3B. Intracellular bacterial survival did not associate to any *Enterobacter* species tested, the presence of colistin resistance, lipopolysaccharide modifications, or virulence in the *Galleria mellonella* infection model. Moreover, intracellular infection caused minimal cytotoxicity in macrophages without evidence of macrophage proinflammatory cell death by pyroptosis.

*Implications of all the available evidence:* Our findings underscore the capacity of *Enterobacter* species, traditionally viewed as extracellular bacteria, to hideout in macrophages without inducing a significant inflammatory response. These properties may further complicate the treatment of antibiotic-resistant *Enterobacter* infections in susceptible populations such as the elderly and neonates. These findings open a door to the development of host-directed therapeutics to enhance bacterial clearance by macrophage-mediated killing.

## Introduction

The *Enterobacter* genus comprises Gram-negative, facultative anaerobic bacteria, belonging to the Enterobacteriaceae family. The taxonomy of *Enterobacter* has evolved very rapidly in recent years, resulting in the designation of numerous new species; pathogenic species are commonly grouped under the *Enterobacter cloacae* complex (Ecc)^1^. Many *Enterobacter* species ^2,3^ are found in the human gut microbiota, but also as pathogens causing bacteraemia, neonatal sepsis ^4-6^, and infections in multiple sites ^2,7,8^. They are also found in aquatic and terrestrial environments, and some species can be symbiotic or pathogenic for plants and insects ^9-12^. Infections are difficult to treat due to multidrug antibiotic resistance (AMR) including resistance to carbapenems and last-resort antibiotics such as polymyxins ^13,14^; for this reason, *Enterobacter* species are included in the WHO’s high priority list of global threat pathogens ^15^ for which new antibiotics are urgently needed.

Despite the abundant information on antibiotic resistance by *Enterobacter* species, much less is known about their infection biology. A handful of studies have reported survival of *E. cloacae* and *E. hormaechei* in human monocytic macrophage cell lines ^16-18^. *E. cloacae* was also reported to form intracellular bacterial communities in uroepithelial cells ^19^, and several *Enterobacter* species can induce apoptosis in human epithelial cells ^20-22^. However, the mechanisms by which *Enterobacter* clinical isolates survive intracellularly have not been explored in any detail. In this study, we have investigated the intracellular trafficking of several antibiotic-resistant *Enterobacter* clinical isolates within human macrophages and determined the macrophage cellular response to the infection using two prototypic species: *E. cloacae* ATCC13047 and *E. bugandensis* 104107. Our data demonstrate that *Enterobacter* strains from several different species can survive in human macrophages for at least 44 hours with no replication. Bacterial survival occurs in a modified late phagolysosomal compartment with reduced acidity and does not induce overt cytotoxicity but primes the macrophages to undergo pyroptosis if a second signal is encountered. These findings underscore the capacity of *Enterobacter* clinical isolates, traditionally viewed as extracellular bacteria, to establish a protective niche in macrophages and possibly delay detection by the innate immune system. This may further complicate the treatment of infection in susceptible populations.

## Methods

### Study design

This work was an exploratory study analysing the intracellular survival of several antibiotic-resistant *Enterobacter* clinical isolates in human cultured and primary macrophages and assessing whether the intracellular infection induces cytotoxicity and pro-inflammatory responses.

### Bacterial strains and culture conditions

The *Enterobacter* clinical isolates used in this study were originally collected by the British Society of Antimicrobial Chemotherapy (BSAC) Surveillance Programmes ^23^ and investigated as part of a case control comparison of colistin-resistant and -susceptible isolates representative of various *Enterobacter* species ^24^. For this study, we used a subset of 11 isolates of clinical origin that included *E. bugandensis* (2 isolates), *E. kobei* (2 isolates), *E. xiangfangensis* (2 isolates), *E. roggenkampii* (1 isolate), *E. hoffmannii* (1 isolate), *E. ludwigii* (2 isolates), and the *E. cloacae* type strain ATCC13047. Full details of bacterial growth conditions, antibiotic susceptibility and the taxonomic assignment of these isolates are described in the appendix (p. 3).

### *Galleria mellonella* Infection

*Galleria mellonella* larvae (UK Waxworms ltd.) weighing between 250 and 300 mg were inoculated with cultures grown to mid-log phase (OD_600_ 0.6). Cultures were adjusted to an OD_600_ of 0.001 (equivalent to 10^5^ CFU/ml) in phosphate-buffered saline (PBS). Ten larvae were infected with each isolate; infections were repeated three times. A control group of larvae was injected with the same volume of PBS. The survival ratio was measured daily for 5 days. More details are given in the appendix (p. 3).

### Lipid A mass spectrometry

Matrix-assisted laser desorption/ionization-time of flight (MALDI-TOF) mass spectrometry (MS) was used to characterise the lipid A modifications present in the LPS of strains with and without challenge with polymyxin B (a colistin analogue). Details of lipid A extraction and analysis are provided in the appendix (p. 3).

### Cell culture, macrophage infection and confocal imaging

THP-1 monocytes (American Type Culture Collection TIB-202) were maintained in RPMI 1640 medium (ThermoFisher Scientific) plus 10% Fetal Bovine Serum (ThermoFisher Scientific). Cells were seeded at a density of 2 × 10^5^ cells/mL and differentiated into macrophages in 80 ng/mL Phorbol 12-myristate 13-acetate (PMA; Sigma-Aldrich) for 3 days. Human monocyte-derived macrophages (HMDMs) were isolated from buffy coats obtained from the Northern Ireland Blood Transfusion Service (Project Reference number 2019/09) according to ethical approval by the Ethics Committee of the Faculty of Medicine, Health and Life Sciences, Queen’s University Belfast (Reference MLHS 19_22). Detailed information on the isolation of HMDMs may be found in the appendix (pp. 3-4). THP-1 and HMDMs cells were infected as described in the appendix (p. 4) with the clinical isolates at various multiplicities of infection (MOI) (indicated in the Figure Legends). After 30 min, infected macrophages were treated for 30 min with 100 μg/mL kanamycin to kill extracellular bacteria (see appendix, pp. 3-4) and then incubated with 50 μg/ml kanamycin for the remaining duration of the experiments. Macrophages were either lysed to enumerate bacteria or processed for imaging using live and immunofluorescence staining microscopy, as applicable (see details in appendix, p.4).

### Cytotoxicity assay

The cytotoxicity of the bacterial infections in macrophages was investigated using the Roche LDH assay kit with a minimum of 2 technical repeats per experiment and up to 6 biological repeats. A detailed protocol is provided in the appendix, p. 4.

### Immunoblotting

Experiments were performed at 5-hours post-infection using a detailed protocol described in the appendix, p.4). Positive controls included LPS-treated (1 μg/mL) macrophages with and without the addition of nigericin (20 μM) to induce pyroptosis and inflammasome priming, respectively. Priming effects were also investigated by exposing the macrophages to heat-killed bacteria with and without nigericin, as before. Blots were treated with primary and secondary antibodies to assess cleavage of caspase-1, gasdermin D (GSDMD) and interleukin-1β (IL-1β) indicating pyroptosis. Detection was performed by incubation with the secondary goat anti-rabbit IRDye 800 cW (LI-COR) antibody and membranes were imaged using the LI-COR Odyssey scanner (see more details in the appendix, p.5).

### Statistical analysis

Since CFU do not follow a normal distribution, we analysed the natural log-transformed values using geometric means. For comparison of two groups in microscopy analysis of infected tissue sections that followed a normal distribution we used a t test. For comparison of multiple groups, a one-way ANOVA was done, with Tukey’s correction for multiple comparisons. p values were considered significant if they were less than 0·05.

## Results

Initial experiments to investigate the interaction between clinical *Enterobacter* isolates and human macrophages employed *E. bugandensis* 104107, a colistin-resistant isolate obtained from respiratory secretions ^24,25^. THP-1 macrophages were infected with 104107 expressing the mCherry red-fluorescent protein. Confocal fluorescence microscopy at early times post-infection, ranging from 15 to 60 min, revealed that the bacteria traffic into *Enterobacter-*containing vacuoles (EcVs) colocalizing with early endosomal antigen 1 (EEA1), an early phagosomal marker **(Figure 1A and appendix, p.7)**. The percentage of EcVs-EEA1 colocalization decreased from 70% at 15- and 30 min post-infection to less than 10% at 60 min **(Figure 1B)**. Transient EEA1 recruitment into early phagosomes ^26^ suggested that intracellular bacteria do not interfere with the initial maturation of EcVs. We next investigated whether EcVs recruit the late phagosomal marker LAMP1. Contrary to the results with EEA1, the percentage of EcVs-LAMP1 colocalization increased over time **(Figure 1A and B, and appendix, pp. 8-9)**, consistent with traffic of the EcV from an early to a late phagosome. Similar results were obtained in THP-1 cells stably expressing GFP-tagged LAMP2 **(Appendix, p. 10)**, also a late phagosomal marker; this experiment revealed that intact bacteria could remain in late phagolysosomes for up to 24-hours post-infection.

**Figure 1:**
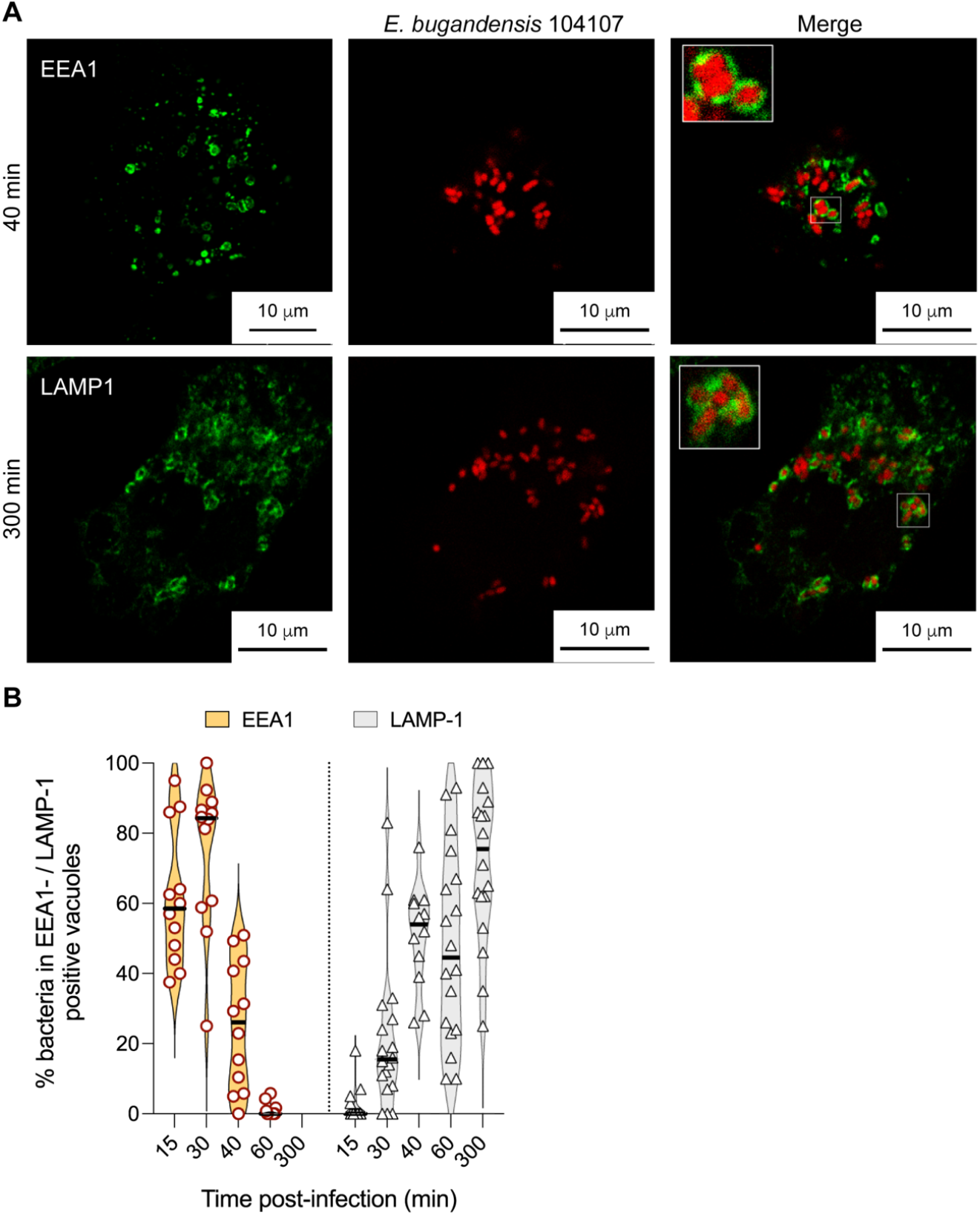
Intracellular trafficking of *E. bugandensis* 104107 from early to late phagolysosomes in THP1 macrophages by confocal immunofluorescence microscopy. (A) Colocalization of live *E. bugandensis* 104107 (red fluorescence) in membrane vacuoles labelled for the early phagosomal marker EEA1 (green fluorescence) at 40 min post-infection, and vacuoles labelled for the late phagolysosomal marker LAMP1 (green fluorescence) at 300 min post-infection (Additional images at various timepoints post-infection are shown in the appendix, pp. 8-10). All images were taken with x 63 magnification on a Leica SP8 confocal microscope. MOI = 120. (B) Graph indicating the percent of intracellular bacteria in EEA1- and LAMP1-positive vacuoles over time. Data were obtained by counting the number of vacuoles with bacteria and the corresponding vacuolar markers on at least 40 macrophages per timepoint over three biological replicates.

The observations in THP-1 macrophages were recapitulated in primary human monocyte-derived macrophages (HMDMs) obtained from buffy coats and differentiated with GM-CSF. *E. bugandensis* 104107-infected HMDMs were followed 3, 5 and 24 h. The results indicated that intracellular bacteria remain confined to LAMP1-positive vacuoles **(appendix, p. 11)**. The infection of HMDMs obtained after stimulation with M-CSF **(appendix, p11)** which induces macrophage polarization into anti-inflammatory phenotypes ^27^, demonstrated that the bacteria also remain within LAMP1 compartments up to 44-hours post-infection. Therefore, we conclude that *E. bugandensis* 104107 can also infect primary human macrophages, residing in a late phagosomal compartment for a relatively long time, and bacterial survival occurs in both pro- and anti-inflammatory macrophages.

Additional live microscopy of infected macrophages using with the vital dye calcein blue, which becomes membrane-impermeable upon entering the cells but stains the cytosolic compartment ^28^, revealed that bacteria remain within vacuoles (**Figure 2)**. Tri-dimensional reconstruction of multiple Z-stacked images of single cells, confirmed that internalised bacteria are located within vacuoles (**appendix, p. 12)**. Some intracellular bacteria, like *Burkholderia cenocepacia*, acquire autophagy markers early after engulfment by macrophages, as demonstrated by the colocalization of the essential autophagy protein LC3B within the bacteria-containing vacuole ^29^. Therefore, we assessed whether LC3B was associated with EcVs. Infected THP-1 macrophages were imaged for LC3B at 3.5-hours post-infection. As a control, we treated THP-1 cells with 70 μM chloroquine overnight, which inhibits autophagy by interfering with the fusion of autophagosomes with lysosomes ^30^. The results demonstrated that 104107 bacterial cells did not colocalize with LC3B (**Figure 2)**, since only diffuse fluorescent LC3B puncta were observed throughout infection, but not in association with fluorescent bacteria. Similarly, autophagosomes were detected in the chloroquine-treated cells as bright fluorescent areas of dense LC3B accumulation, but none of these contained bacteria (**Figure 2)**. In contrast, LC3B was associated with *B. cenocepacia-*containing vacuoles (**appendix, p. 13)**. Together, these experiments exclude the possibility that EcVs are autophagosomes.

**Figure 2:**
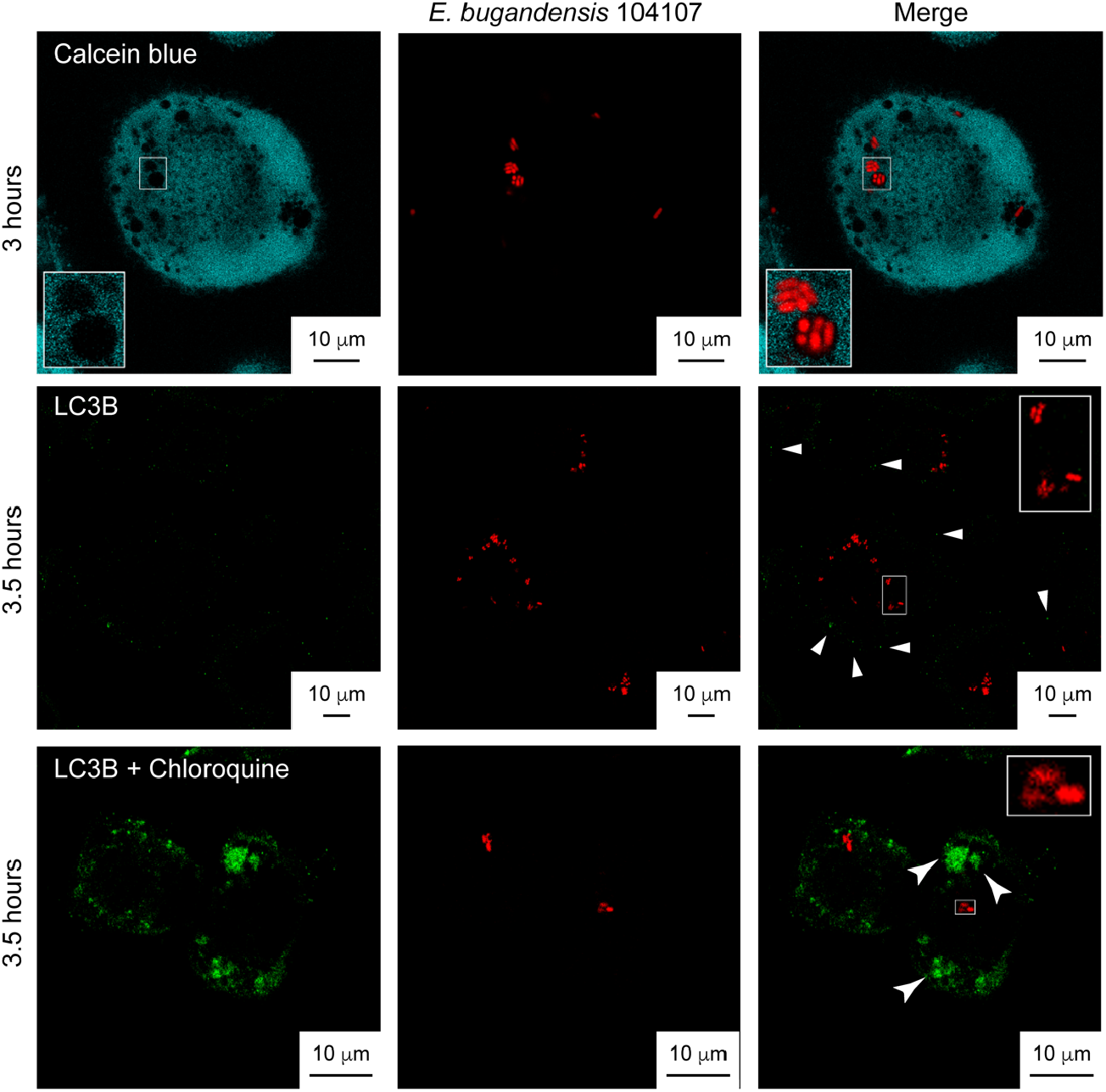
Intracellular *E. bugandensis* 104107 remain within membrane vacuoles and do not traffic to autophagosomes. (Top panel) Live cell imaging of THP-1 macrophages infected with *E. bugandensis* 104107 (red fluorescence) at 3 h post-infection and stained with calcein blue (original blue fluorescence was pseudocoloured as cyan for better contrast), which does not enter membrane compartments. Images taken on a Leica Stellaris-5 confocal microscope with x 100 magnification. MOI = 15. Additional images at 7-hours post-infection and a 3D reconstruction model of an infected calcein blue-stained macrophage cell is shown in the appendix p. 12). (Middle panel) Immunofluorescent microscopy images of infected macrophages (red fluorescence) stained for the autophagy marker LC3B at 3.5 h post-infection. Arrows in the merged image indicate LC3B puncta (green fluorescence) that do not colocalize with the intracellular bacteria. (Lower panel) similar experiment as in the panel above with macrophages treated with 70 μM chloroquine to inhibit autophagy, which shows accumulation of the LC3B marker (green fluorescence) in larger clumps (arrows) but without colocalization with the intracellular bacteria (red fluorescence). Images in middle and lower panels were taken with x 63 magnification on a Leica SP8 confocal microscope. MOI = 120.

To better define the EcVs, we investigated the 104107 colocalization with dextran-TMR, a fluid phase marker that traffics to lysosomes. THP-1 macrophages pre-labelled with dextran were infected with 104107 for 2 h. Confocal microscopy revealed that bacteria colocalize with dextran-rich compartments **(Figure 3A)** presumed to be lysosomes. One other characteristic of lysosomes is their low pH. We estimated the intraluminal pH of the EcVs using lysotracker green DND-26 in infected THP-1 cells. Only ∼25% intracellular 104107 bacteria were found in bright lysotracker-positive compartments (**Figure 3B and C**). Compared to bacteria that did not colocalise with lysotracker, the bacteria present in the strongly acidic vacuoles appeared weakly labelled and, in some cases, the entire vacuolar lumen was red-fluorescent **(Figure 3B and C)**, suggesting that low pH compromised the bacterial cell envelope integrity ^31^. As expected, no colocalization with lysotracker was observed in cells treated with the vacuolar proton-ATPase inhibitor bafilomycin (**Figure 3D**). Together, these experiments demonstrate that engulfed 104107 mainly traffic into a modified *Enterobacter*-containing late phagosomal compartment that delays acidification.

**Figure 3:**
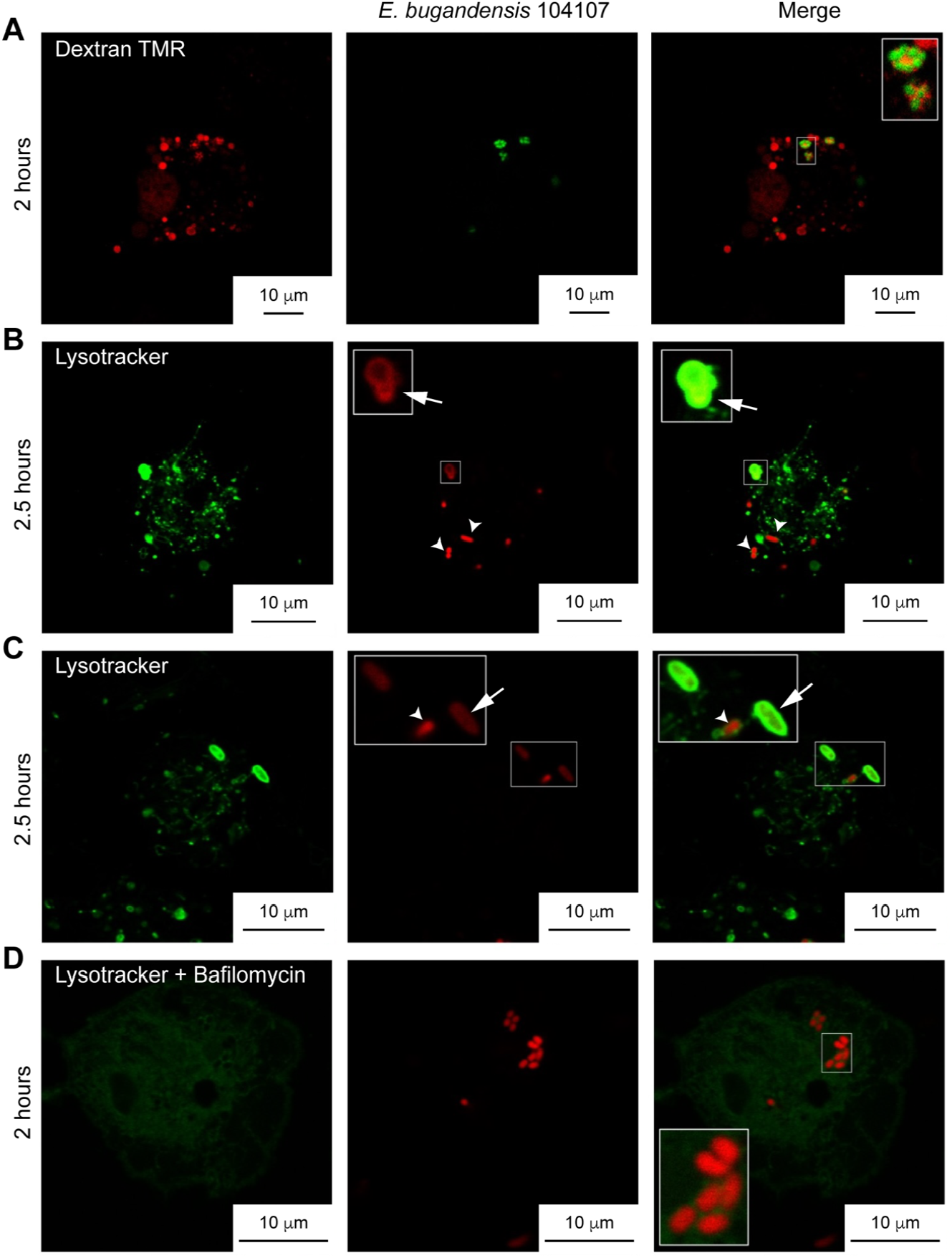
Intracellular *E. bugandensis* 104107 can traffic to a phagolysosome that delays acidification. (A) THP1 macrophages were pre-treated before infection with the fluid phase marker dextran TMR to preload phagolysosomes with red-fluorescent particles. Cells were infected with *E. bugandensis* 104107 containing pLS2 (encoding GFP; green bacteria) and imaged 2 h post-infection at x 100 magnification on an SP8 confocal microscope. MOI = 50. (B) THP-1 macrophages infected with *E. bugandensis* 104107 expressing mCherry (red fluorescence) were treated with the fluid phase marker Lysotracker green DND and imaged at 2.5 h post-infection. Arrows indicate bright, acidic vacuoles where the entire vacuolar lumen was red-fluorescent. Arrowheads point to intact bacteria that did not colocalise with lysotracker. (C) Another field of view from the same experiment in B showing weakly red-fluorescent bacteria that colocalise with Lysotracker-stained vacuoles (arrows), while bright red bacteria do not (arrowheads). (D) Infected THP1 macrophages treated with bafilomycin to inhibit phagosomal acidification did not show bacterial colocalization with Lysotracker at 2 h post-infection; all bacteria appear to be intact and display similar levels of red fluorescence. For experiments shown in panels B-C, live fluorescent images were captured using a Leica Stellaris-5 confocal microscope, x 100 magnification. MOI = 15.

The bacterial load in infected macrophages was quantified by treating extracellular bacteria with kanamycin throughout the course of infection. While bacteria were not detected in the medium, macrophage cell lysates upon detergent treatment at 2- and 5-hours post-infection revealed a similar bacterial load as the inoculum at time 0 (**Figure 4A)**. The lysate at 24-hours post-infection showed a 3-fold reduction in bacterial counts. A small number of bacterial counts was detected in the last wash prior to detergent-mediated cell lysis in all sampling times, which we attributed to a low level of spontaneous macrophage lysis (**Figure 4A)**. Overall, the recovery of intracellular bacteria over time at similar loads as the initial inoculum demonstrates that the bacteria remain viable but do not replicate. This conclusion was further validated by fluorescence dilution experiments involving the fluorescent peptidoglycan precursor D-amino acid 7-hydroxycoumarincarbonylamino-D-alanine (HADA) ^32^. Fluorescent bacteria could be detected in LAMP1-labelled vacuoles when macrophages were infected for 6 h with bacteria grown with HADA (**Figure 4B)**. This suggests that new cell wall synthesis, and hence bacterial cell division, did not occur, consistent with the absence of bacterial replication. In contrast, HADA-labelled bacteria growing in culture medium lost blue fluorescence by 3 h, in contrast to the bacteria treated with the bacteriostatic antibiotic chloramphenicol **(Figure 4C)**, which retained fluorescence. Combined, these experiments confirm that 104107 remains viable and persists in human macrophages without replication.

**Figure 4:**
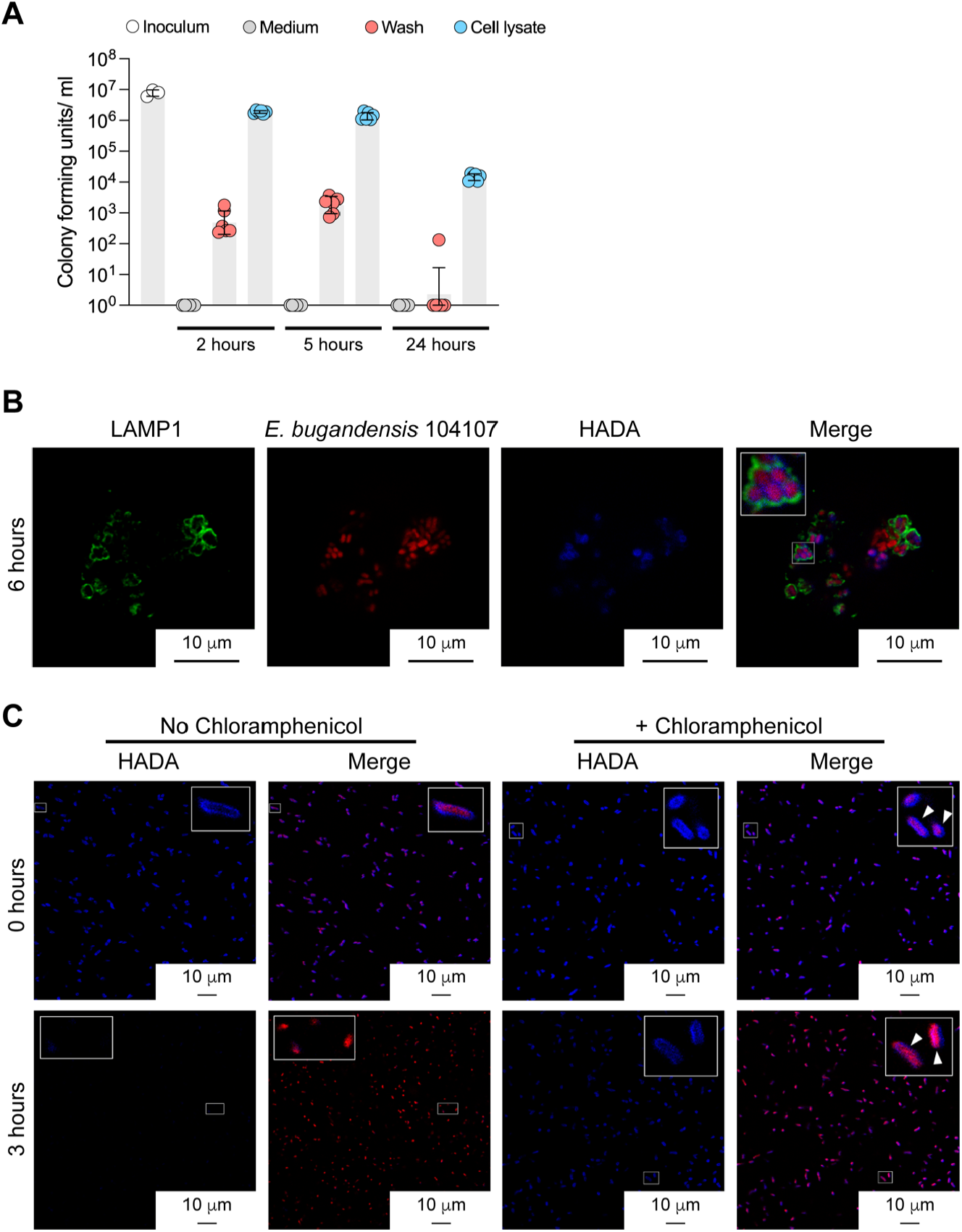
*E. bugandensis* 104107 bacteria do not replicate in THP1 macrophages. (A) Intracellular bacteria enumeration (CFU/ml) at various times post-infection comparing the number of bacteria recovered at each sampling timepoint from the medium (antibiotic control), last wash prior macrophage lysis (macrophage intactness control), and cell lysate with the inoculum at time 0. After 1-hour post-infection, extracellular bacteria were killed by adding 100 μg/mL kanamycin for 30 min and then 50 μg/mL remained in the medium for the duration of the experiment. (B) Bacteria were pretreated with HADA (blue fluorescence) prior to infection of THP1 macrophages and imaging by immunofluorescence confocal microscopy at 3 and 6 h post-infection. Merged image shows HADA-positive bacteria (red fluorescent bacteria surrounded by blue fluorescence halos colocalizing with LAMP1. Images were taken with x 100 magnification on a Stellaris-5 confocal microscope. MOI = 15. (C) Live *E. bugandensis* 104107 were pretreated with HADA and with or without 50 μg/ml chloramphenicol to prevent replication and imaged live at time 0 (top panel) and 3 h (lower panel). Arrows in the merge images indicate HADA-labelled bacteria (blue halos surrounding bacterial cells) a time 0 (top panel), and the absence of HAD labelling after 3 h of incubation at 37 °C (lower panel).

Next, we investigated whether the observations with *E. bugandensis* 104107’s intracellular survival could be extended to other *Enterobacter* species. We performed THP-1 infections using mCherry-fluorescent derivatives of antimicrobial resistant Ecc clinical isolates with different levels of colistin resistance, which were characterised in a previous study ^24^. All these isolates colocalized with LAMP1-positive compartments at 3.5-hours post-infection (**Figure 5A)**, and 9 of them infected ∼60% of macrophages (**Figure 5B)**. We conclude that Ecc bacteria from different species traffic into late phagolysosomes, including the ATCC13047 *E. cloacae* type strain. Notably, all the infecting bacteria induced low levels of macrophage cytotoxicity (ranging from 2 to 7%) in comparison with the 15% cytotoxicity of lipopolysaccharide-treated macrophages (**Figure 5C)**, and despite some minor differences in toxicity in some strains when infections with live vs. heat-killed (HK) bacteria were compared. The intracellular survival of the clinical isolates also did not correlate with their level colistin/polymyxin resistance, their lipopolysaccharide lipid A structures, or their levels of relative virulence in the *Galleria mellonella* infection model (**Figure 5D and appendix, pp. 14-16)**. Together, these experiments demonstrate that Ecc clinical isolates survive intracellularly in human macrophages without inducing high levels of cytotoxicity.

**Figure 5:**
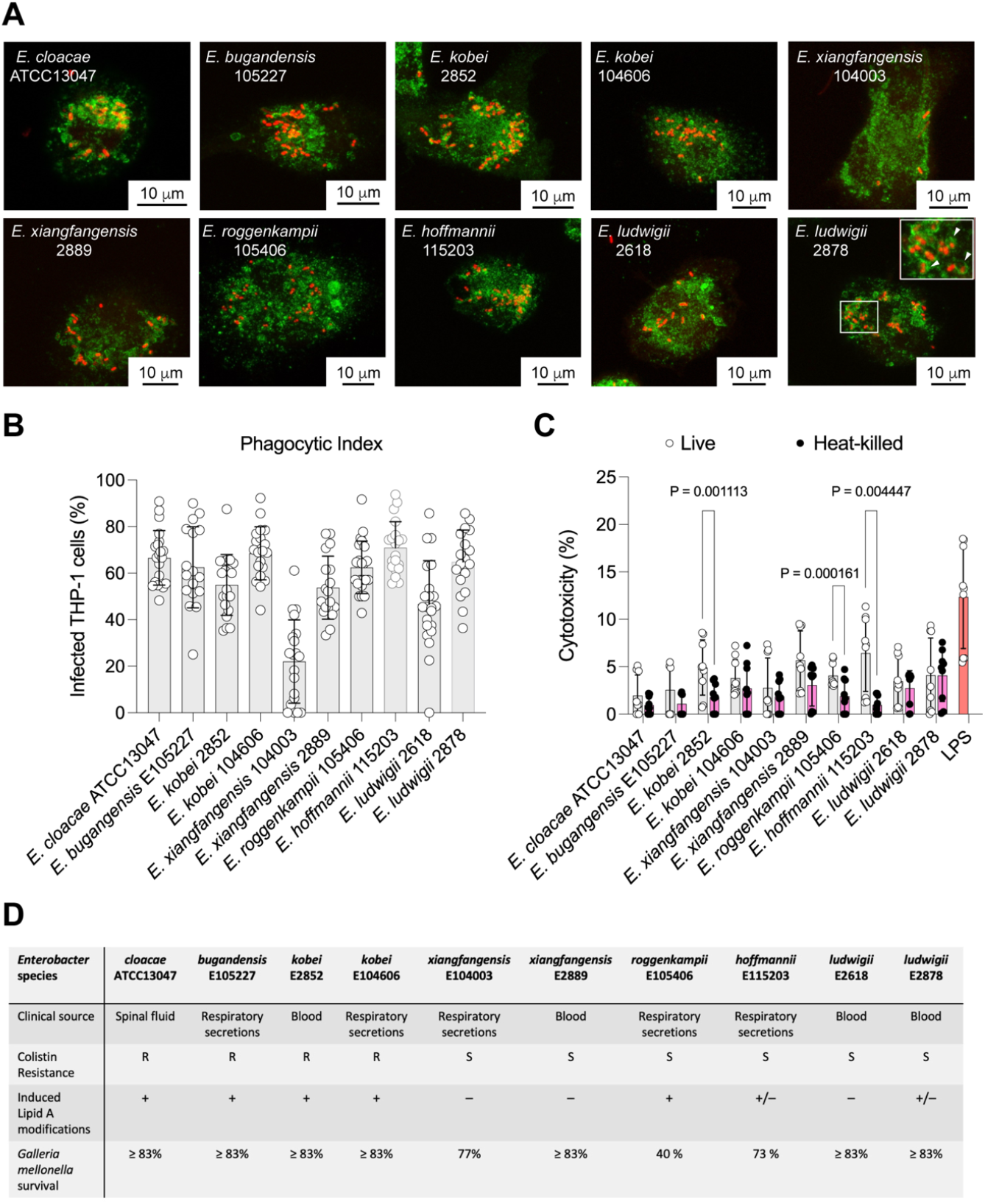
Ecc clinical isolates traffic to LAMP1-positive compartments in THP1 macrophages causing minimal cytotoxicity. (A) THP-1 macrophages infected with live Ecc isolates (all expressing red fluorescent mCherry) at 3.5 h post-infection colocalize with LAMP1 (green fluorescence). Images taken at X 63 magnification on Leica SP8 confocal microscope. MOI = 120. (B) The percentage of infected macrophages for each isolate was assessed by counting an average of 40 fields of view. (C) Infected macrophages with the live and heat-killed Ecc isolates display minimal cytotoxicity at 5 h posit infection, which was measured by LDH release. (D) Characteristics of the Ecc isolates comparing their clinical source, colistin resistance data (published in reference ^24^), polymyxin-induced lipid A modifications (details are in the appendix, pp. 14-15), and pathogenicity in the *Galleria mellonella* infection model (see appendix, p. 16).

One of the most common outcomes of bacterial infection in macrophages is the elicitation of a proinflammatory form of programmed cell death known as pyroptosis ^33,34^. Typically, pyroptosis involves the activation and self-cleavage of pro-caspase-1, which in turns cleaves Gasdermin D (GSDMD) and pro-interleukin-1β (IL-1β). The N-terminal domain of GSDMD ((GSDMDNT) oligomerises and forms a pore in the cell membrane leading to the escape of mature IL-1β and cell lysis ^35^. To investigate pyroptosis during Ecc infection, THP-1 macrophages were infected with live *E. cloacae* ATCC13047 and *E. bugandensis* 104107; western blotting was used to assess the levels of caspase-1 and GSDMD cleavage, and IL-1β secretion in cell lysates and supernatants at 5-hours post-infection. This timepoint was selected to ensure all intracellular bacteria have completed their trafficking into phagolysosomes. As controls, macrophages were treated with lipopolysaccharide (LPS) only, LPS plus nigericin (positive control for pyroptosis via the activation of the NLRP3 inflammasome ^36^), and live or HK bacteria also in the presence of nigericin. In the experiments with ATCC13047 (**Figure 6A)**, the active p20 caspase-1 form was clearly detected in supernatants of nigericin-treated macrophages. The corresponding cell lysates of these samples revealed 2-fold less amount of procaspase-1 (∼45 kDa), as determined by densitometry, than macrophages exposed to LPS-only or infected with live bacteria. Blots of cell lysates probed for GSDMD revealed the presence of the ∼31 kDa GSDMD N-terminal fragment (GSDMDNT) in all cases. However, the amount of this product was 4-fold higher in nigericin-treated cells (**Figure 6A)**. GSDMDNT, which upon GSDMD cleavage oligomerises and inserts in the cell membrane ^37,38^, was also detected in cell supernatants of nigericin treated cells, suggesting that cell lysis had occurred in these samples. The IL-1β blots revealed a similar pattern showing that the lysates of nigericin-treated macrophages exposed to LPS or infected with live ATCC13047 contained 2-fold and 10-fold less pro-IL-1β, respectively than the nigericin-untreated macrophages (**Figure 6A**). Both IL-1β and pro IL-1β were also detected in the supernatants of nigericin-treated cells. Together, the data suggest that cell lysis due to pyroptosis occurs only in the nigericin-treated cells but not in cells infected with live ATCC13047 bacteria. Similar results were recapitulated using *E. bugandensis* 104107 (**Figure 6B**). In this case, we also included macrophages incubated with live and HK bacteria, both treated and untreated with nigericin. Densitometric analyses indicated similar patterns as shown with *E. cloacae* ATCC13047 infections. The only exception was the detection of IL-1β in the supernatant of nigericin untreated macrophages infected with live 104107 **(Figure 6B)**. Since p20 caspase-1 and GSDMDNT were not detected, we cannot conclude that 104107 induced pyroptosis, suggesting that the secretion of IL-1β could be due to GSDMD-independent mechanisms ^39^. However, this observation was unique to this strain since processed IL-1β was not detected in supernatants from macrophages infected with ATCC13047 and *E. hoffmannii* 115203 (**Figure 6A and Appendix, p. 17**). Moreover, detection of lactate dehydrogenase in the supernatants of macrophages infected with live ATCC13047 or 104107, as a proxy for macrophage cell lysis, show very low levels of cell lysis (≤ 15%) compared with ∼80% lysis of the macrophages treated with LPS and nigericin (**Figure 6C**). These results demonstrate that macrophages infected with clinical isolates of three different *Enterobacter* species do not undergo pyroptosis, supporting the notion that engulfed *Enterobacter* species survive intracellularly in macrophages without inducing significant cytotoxicity.

**Figure 6:**
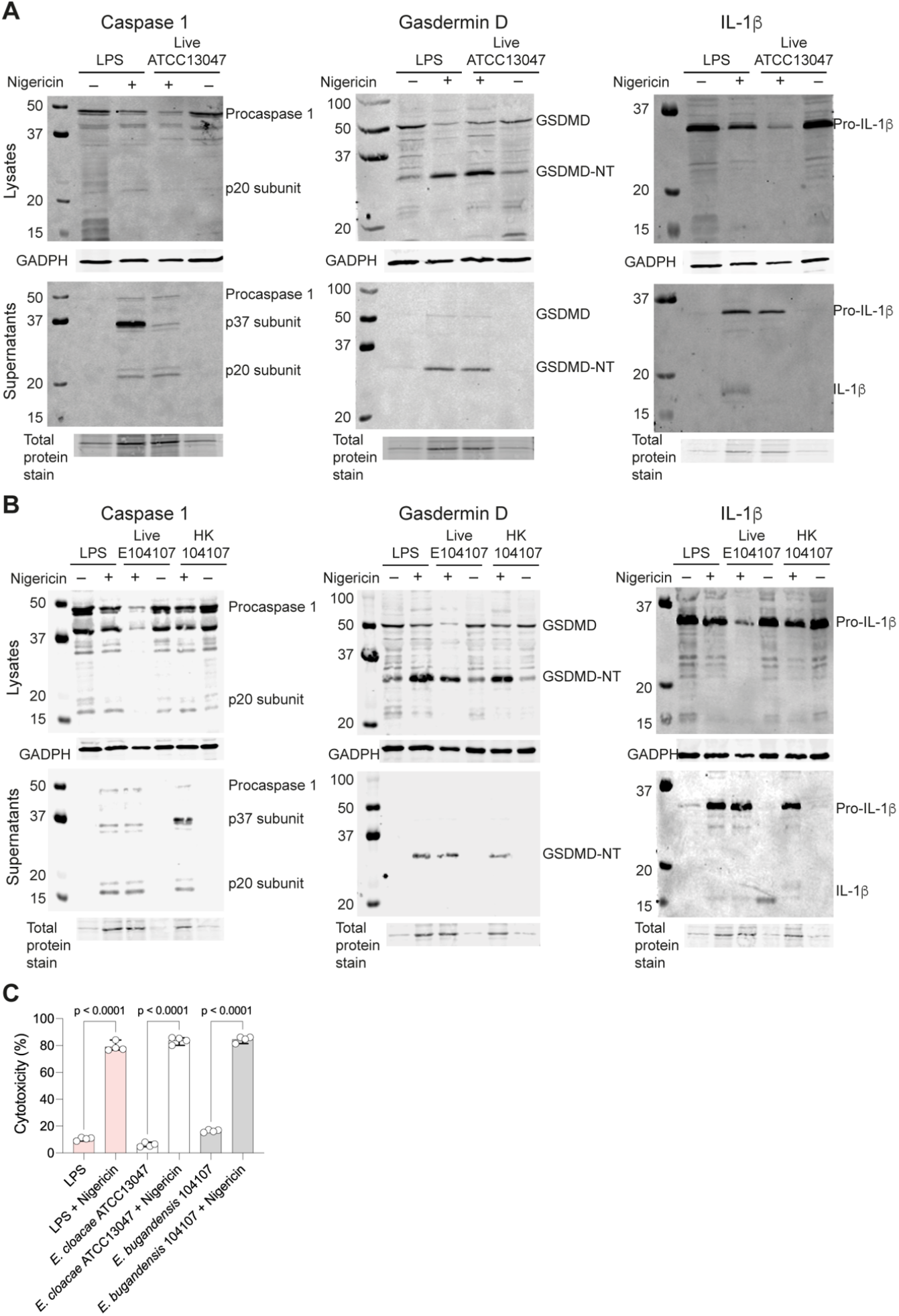
Macrophage responses to *E. cloacae* ATCC13047 and *E. bugandensis* 104107 infections. Western blots showing full length and cleaved caspase-1, gasdermin D (GSDMD), and IL-1β in lysates and supernatants of live *E. cloacae* ATCC13047-infected THP1 macrophages (A), and live and heat-killed (HK) *E. bugandensis* 104107-infected THP1 cells. Both sets of experiments were done by triplicate using cells at 5 h post-infection. MOI = 40. For inflammasome priming and pyroptosis controls, uninfected macrophages were treated with and 1 μg/mL LPS and LPS + 20 μM Nigericin, respectively. (C) Percent cytotoxicity from LDH assays in culture supernatants of infected macrophages treated and untreated with Nigericin and the LPS- and LPS + Nigericin controls at 5 hours post-infection.

## Discussion

This study discovered that clinical isolates of different *Enterobacter* species can survive intracellularly while remaining viable in human macrophages. Survival occurs in bacteria-containing vacuoles resembling late phagolysosomes that delay acidification, suggesting the bacteria survive in a modified vacuole that delays acidification. Two key features separate *Enterobacter* from other known facultative intracellular bacteria. First, experiments directly measuring the intracellular bacterial load after killing extracellular bacteria, or examining the dynamics of wall synthesis in the intracellular bacteria, demonstrate that engulfed bacteria do not replicate intracellularly. This is demonstrated by similar numbers of recovered bacteria from infected macrophages over time and the initial inoculum at time 0. However, bacteria remain intact and viable since they express the endogenous mCherry red-fluorescent protein encoded by a plasmid at all timepoints investigated extending up to 44 hours in HMDMs. Moreover, bacteria can be recovered at 24-hours post-infection by enumeration of CFUs, while no bacteria can be recovered from the culture medium treated with kanamycin, an antibiotic that cannot efficiently cross eukaryotic cell membranes. These observations imply that *Enterobacter* isolates can hide within macrophages as viable, nonreplicating bacteria, joining a long list of bacterial pathogens that can persist in macrophages ^40^. We posit that the ability of *Enterobacter* isolates to survive in macrophages provides nonreplicating bacteria with an additional strategy to escape antibiotics. This may be clinically relevant, since nonreplicating intracellular *Enterobacter* may escape the action of antibiotics that can penetrate the cell membrane of eukaryotic cells, such as carbapenems, tetracyclines and fluroquinolones. Moreover, infected macrophages could act as Trojan horses to disseminate *Enterobacter*. This idea is reinforced by a study demonstrating the association of *E. hormaechei* with atheromatous tissues and its isolation by co-cultivation of plaque tissue homogenates with THP-1 macrophages ^17^. Also, certain clinical strains of *E. hormaechei* associated with neonatal sepsis were shown to persist in U937 human macrophages and invade rat brain capillary endothelial cells ^16^.

The second salient and surprising feature of *Enterobacter*-macrophage infection is that prolonged bacterial intracellular survival results in minimal macrophage cytotoxicity, suggesting low pathogenicity of *Enterobacter* species. Persistence depends on how well the pathogen can survive in the host, and the balance between the cost of infection versus the costs and effectiveness of the immune response against the infection ^41^. Our results agree with the notion that while the magnitude of pathogenicity drives virulence, it does not necessarily increases the rate of host clearance, allowing for persisting bacterial populations ^41^. Indeed, we show that survival in macrophages does not correlate with the pathogenicity of our clinical isolates in the *Galleria mellonella* infection model. Moreover, survival cannot be explained by resistance to antimicrobial peptides since both colistin-resistant and colicin-sensitive isolates can survive in macrophages, or with the lipid A modifications associated with resistance to cationic antimicrobial peptides. The lack of correlation of intracellular survival and colistin resistance further suggests that intracellular bacteria may be able to resist antimicrobial peptides. The intracellular *Enterobacter* isolates investigated here do not cause any significant cell lysis and do not elicit pyroptosis, suggesting that intracellular *Enterobacter* could manipulate cell death pathways. In primary human monocytic macrophages, differentiation into proinflammatory or anti-inflammatory types using GM-CSF or M-CSF, respectively, do not affect the intracellular fate of *Enterobacter*. While the mechanism of intracellular survival in macrophages without an overt proinflammatory response remains to be elucidated, it may be possible that intracellular bacteria contribute to remodelling the macrophages’ cell metabolism. Metabolic reprogramming affecting mitochondrial metabolism and antagonizing programmed cell death by apoptosis to facilitate bacterial persistence in bladder cells has been reported in uropathogenic *Escherichia coli* ^42^. *Enterobacter* species form intraepithelial communities in bladder cells, which might provide a reservoir for symptomatic and asymptomatic urinary infections ^19^. Metabolic reprogramming in macrophages infected with *Klebsiella pneumoniae*, a pathogen closely related to *Enterobacter*, has been shown to modulate cell death pathways in a bacterial type VI secretion system (T6SS)-dependent manner ^43^. A screen for *in vivo* fitness-associated genes in *E. cloacae* ATCC13047 upon infection in *Galleria mellonella* discovered several metabolic genes, genes encoding transcriptional regulators, surface protein- and T6SS-encoding genes ^44^. However, it is unclear if T6SS effectors play a direct role in survival since another *E. cloacae* isolate does not appear to require T6SS for invasion and proliferation within host cells ^18^.

In summary, our findings underscore the capacity of AMR *Enterobacter* species, traditionally viewed as extracellular bacteria, to hideout in macrophages without significantly alerting the innate immune system. We posit our observations have clinical implications since intracellular survival of nonreplicating *Enterobacter* bacteria may further complicate the treatment of *Enterobacter* infections in susceptible individuals.

## Contributors

GP, HJP, and MAV wrote the manuscript. GP, HJP, AJGA, FB, IGR, MG, SOK, and HM contributed to manuscript. GP, HJP, FB, and HM performed imaging experiments. AJGA developed the kanamycin killing assay to enumerate intracellular bacteria. IGR, MG, and SOK performed in-vivo *Galleria* infections. IGR and MG characterised the antibiotic resistance of the clinical isolates. IGR analysed the lipid A profiles by mass spectrometry. MAV performed the in-silico taxonomy of the isolates and some of the statistical analysis. MAV supervised GP, HJP, AJGA, FB, IGR, MG, SOK, HM and verified the data. All authors had full access to all the data in the study and had final responsibility for the decision to submit for publication.

## Acknowledgments

We thank Dr. Shazad Mushtaq, Antimicrobial Resistance and Healthcare Associate Infections Reference Unit, UK Health Security Agency, for the gift of the clinical *Enterobacter* strains used in this research, Dr. Ileana Micu and Dr. Ryan Delaney for technical assistance with confocal imaging and 3D reconstructions, and Karolina Woljtania for technical assistance with the HADA experiments. GP, HJP, AJGA, and FB were supported by PhD studentships from the Department of Economy of Northern Ireland. SOK was supported by an Internship from the International Association for the Exchange of Students for Technical Experience.

