## Appendix for "Clinical isolates of *Enterobacter* species can persist in human macrophages without bacterial replication and minimal cellular cytotoxicity"

**SUPPLEMENTARY APPENDIX**

This appendix is part of the original submission by Parau G, Parks HJ, Anderson AJG et al. Clinical isolates of *Enterobacter* species can persist in human macrophages without bacterial replication and minimal cellular cytotoxicity

### Table of Contents

Supplementary Materials and Methods — Page 3

Table S1 - Antibodies and Fluorescent Reagents — Page 6

Supplementary Results

*E. bugandensis* 104107 trafficking to late phagolysosomes in THP-1 and HMDM macrophages

Figure S1 — Page 7

Figure S2 — Page 8

Figure S3 — Page 9

Figure S4 — Page 10

Intracellular *E. bugandensis* 104107 remains in vacuoles that are not autophagosomes

Figure S5 — Page 11

Figure S6 — Page 12

Lipid A modifications of *Enterobacter* clinical isolates upon challenge with polymyxin B

Figure S7 — Page 13

Table S2 — Page 14

*Enterobacter* clinical isolates display varying levels of virulence in *Galleria mellonella*

Figure S8 — Page 15

Intracellular *E. hoffmannii* 115203 does induce Pro-IL-1β cleavage and release of mature IL-1β into macrophage cell supernatants

Figure S9 — Page 16

Supplementary References — Page 17

### Supplementary Materials and Methods

#### Bacteria and plasmids

*Enterobacter cloacae* complex isolates were collected from bloodstream and lower respiratory infections as part of the extensive BSAC surveillance programme [1,2]. For confocal microscopy experiments these isolates were transformed with plasmids pJT04 or pLS2. Plasmid pJT04 is a modified pDA12 [3], which confers tetracycline resistance and encodes a constitutively expressed mCherry red-fluorescent protein gene. Plasmid pLS2 is a derivative of pML7-eGFP [4] encoding tetracycline resistance. *Burkholderia cenocepacia* isolate DFA35 (*B. cenocepacia* K56-2 Δ*atsR*) [5], carrying pJT04, was used as a control in some of the macrophage infection experiments. Plasmids were introduced in the *Enterobacter* isolates by electroporation [6]; pJT04 was introduced in *B. cenocepacia* DFA35 by triple parental mating [7].

All bacterial strains were routinely grown at 37°C in high-salt Lysogeny medium (LB; Melford) and supplemented, when required, with a final concentration of 50 µg/mL of tetracycline. For experiments designed to examine modifications in the lipid A lipopolysaccharide moiety, bacteria were also grown in cationic-adjusted Mueller-Hinton broth (CAMH) with or without polymyxin B (PMB), as indicated below under Lipid A mass spectrometry. PMB was prepared as a stock solution containing 12 mg/mL of polymyxin B powder (Sigma), 0.2% (w/v) bovine serum albumin (BSA; Sigma) and 0.01% (v/v) acetic acid (Honeywell). Colistin and PMB susceptibility of the isolates was determined by broth microdilution [2], and interpreted using 2019 EUCAST breakpoints for Enterobacterales, which score colistin as susceptible, <2 µg/mL or resistant, >2 µg/mL.

#### Taxonomy of the clinical isolates

The species of the clinical isolates was determined by estimating their genetic relatedness based on whole-genome average nucleotide identity (ANI) measurements using FastANI [8]. Results were refined by digital DNA:DNA hybridisation (dDDH) using the Type Strain Genome Server (TYGS) [9].

#### *Galleria mellonella* Infection

*Galleria mellonella* larvae (UK Waxworms ltd.) were stored at 16°C for up to 3 weeks from delivery. Larvae used for infections weighed between 250-300 mg and appeared healthy based on movements and absence of melanisation. Bacteria were prepared by refreshing overnight LB cultures to mid-log phase, which corresponded to an optical density ($OD_{600}$) of 0.6. Inoculum was adjusted to an $OD_{600}$ of 0.001, which corresponds to ~$10^5$ colony forming units (CFU) per mL, in phosphate-buffered saline (PBS). Infection was carried out using a sterile Hamilton syringe equipped with a hypodermic needle (BD microlance, 30 gauge, 13mm). Ten µL of the bacterial suspension was injected into the rear proleg of each larva. Ten larvae were infected for each isolate and the infections were repeated three times; in addition, a control group was injected with 10 µL of PBS. Larvae survival was scored daily over 5 days.

#### Lipid A mass spectrometry

Matrix-assisted laser desorption/ionization-time of flight (MALDI-TOF) mass spectrometry (MS) was used to characterise lipid A modifications present in the LPS of strains challenged with PMB. The lipid A extraction and analysis was performed as previously described [10,11]. Overnight cultures were refreshed and grown to mid-log phase in CAMH without PMB or supplemented with 1 µg/mL PMB for the sensitive isolates and 10 µg/mL for the resistant isolates. Bacteria were sedimented by centrifugation at 4000 xg for 10 minutes and washed three times in 1 mL of 0.1 M citric acid (Sigma). The pellet was re-suspended in 1 mL of 0.1 M citric acid and boiled at 100 °C for 90 min. The samples were cold down at room temperature and centrifuged 5 minutes at 12000 rpm. After removing the supernatant, the pellet was resuspended in 20 µL of 0.1 M citric acid. To prepare the matrix, 50 µL of 0.1 M citric acid were saturated with 2,5- dihydroxybenzoic acid (DHB; Sigma) and the mix was incubated for 20 minutes in a sonicator bath. The matrix was centrifuged for 5 minutes at 12000 rpm before use. Five µL of the samples were desalted with 5 µL of Dowex 50WX8 (Sigma) converted into its ammonium form. Finally, 1 µL of each sample was loaded onto a different spot of a polished steel target and immediately 1 µL of the matrix was added and mixed with the sample. The target was inserted into a Bruker Autoflex MALDI-TOF mass spectrometer; data acquisition and analysis were performed using the Bruker Flex Analysis software.

#### Isolation of human monocyte-derived macrophages (HMDMs)

HMDMs were obtained from buffy coats provided by the Northern Ireland Blood Transfusion Service (Project Reference number 2019/09) under the ethical approval by the Ethics Committee of the Faculty of Medicine, Health and Life Sciences, Queen's University Belfast (Reference MLHS 19_22). Buffy coats were separated by Ficoll-Paque gradient density fractionation and processed as described [12]. Approximately 15 million cells were plated per

10 cm dish in 15 ml of RPMI medium supplemented with 50 ng/mL GM-CSF. The cells were cultured for 6-7 days, and the medium was replenished on day 5 by adding 5 mL of RPMI + 50 ng/mL GM-CSF per plate.

**Macrophage infection**

PMA-treated THP-1 and HMDM cells were seeded at 2 x $10^5$ cells/mL on 19-mm coverslips in 12-well plates. THP-1 cells were differentiated to adherent macrophages by incubation with PMA (80 ng/mL) for 72 hours and allowed to recover overnight in PMA-free RPMI. Overnight cultures of Ecc isolates (carrying pJT04) were sub-cultured to $OD_{600}$ of 0.05 in 5 mL LB supplemented with 50 µg/mL tetracycline and incubated for 3 hours at 37 °C with shaking at 180 rpm. Log phase Ecc were used for infections. In some experiments using heat-killed bacteria as a control, bacteria were heated at 80 °C for 15 minutes. The strains were washed three times with RPMI prior to infection. Multiplicity of infection (MOI) values were calculated for each strain. The infection was synchronized by centrifugation for 3 minutes at 1000 rpm. Ecc isolates were in contact with the cells for 30 minutes or 1 hour (indicated in the experiments) at 37 °C, after which the cells were washed 3 times with PBS and incubated with fresh medium containing 50 µg/mL kanamycin to inactivate extracellular bacteria. Infections were conducted at various times ranging from 1 hour to 44 hours (indicated in the individual experiments described in this work).

**Quantification of intracellular bacteria**

*E. bugandensis* 104107 was cultured to $OD_{600}$ 0.8 in LB; the bacterial pellet was washed three times in RPMI and adjusted by OD to provide an MOI=15. 2x$10^5$ THP-1 macrophages were infected with bacteria and spun down at 1000 x rpm for 5 minutes to synchronise infection. Cells were incubated with bacteria for 1 hour and then washed thrice with warm RPMI. Kanamycin (100 µg/mL final concentration) was added for 1 hour to kill extracellular bacteria. Thereafter, the concentration of kanamycin was reduced to 50 µg/ml for the duration of infection (2 hours, 5 hours and 24 hours post-infection). At the indicated times, medium was collected bacterial enumeration and cells washed trice with warm RPMI; the last wash was also collected for bacterial enumeration. The cell pellet was lysed with 1% Triton X-100 for 10 min. Medium, third wash, and lysate samples were serially diluted and spotted as 10 µL drops onto LB agar. Plates were incubated overnight at 37°C and CFU/mL was calculated. Infections were performed in duplicate over 3 biological repeats at different days.

**Fluorescence microscopy**

UV-sterilized coverslips were placed in 12-well cell culture plates prior to seeding THP-1 or HMDMs. To prepare samples for microscopic analysis at various times post-infection, infected cells were washed with PBS, fixed with 4 % paraformaldehyde (PFA) for 15-20 minutes and stored in 14 mM ammonium chloride at 4 °C to quench free aldehyde groups, before being washed 3 times in PBS and permeabilised with 0.5 % (w/v) Saponin for 30 minutes at room temperature and then incubated in a humidified chamber at room temperature for 1 hour with the appropriate primary antibodies (see Table S1 in the appendix, p. 6). Antibodies were prepared with 0.5 % Saponin in FBS; 80 µL of antibody/Saponin/BS mixture was used per 19 mm coverslip (8 µL FBS, 72 µL Saponin). The coverslips were incubated for 1 hour with the primary antibodies (antibodies and their working concentration are indicated in Table S2), in a humidifying chamber. After incubation with the secondary goat anti-rabbit AlexaFluor 488 for 1 hour, coverslips were mounted on to glass slides using Fluoroshield with 1,4- Diazabicyclo [2.2.2] octane to reduce bleaching during the image capture. Image acquisition was carried out on Leica SP8, SP5 or stellaris confocal microscopes.

For live imaging, infected cells were seeded directly into 8-well chamber slides (Ibidi GmbH) and incubated with fluid phase markers (see Table S1 on p. 6) at various times pre- or post-infection (depending on the type of marker and experiment). To assess bacterial trafficking into a phagolysosome, cells were incubated with Dextran TMR overnight prior to infection. Vascular pH was estimated by adding Lysotracker 5 minutes at the end of the timepoint before imaging. Calcein blue was also added at the end of infection and incubated for 5 minutes prior to imaging.

To fluorescently stain the bacterial cell wall peptidoglycan, bacterial cultures at log phase were incubated with the peptidoglycan precursor, D-amino acid 7-hydroxycoumarincarbonylamino-D-alanine (HADA) [13], for 1 hour at 37 °C with shaking and then washed in PBS pH 7.0.

Images captured at random were visualised using the Leica application software (LAS X). Zoomed insets were prepared using Fiji v2.0.0-rc 69/1.52p. Imaris 3D V9.4 software (Oxford Instruments) was used for 3D reconstructions and visualisation of Z-stacks.

**Determination of cytotoxicity in macrophages**

The cytotoxic effect of the bacterial infections in macrophages was investigated using the Roche LDH assay kit with a minimum of 2 technical repeats per experiment and up to 6 biological repeats. A high control consisting of cells treated with 1 % Triton X-100 was compared to live and HK infected wells, as well as low control samples from uninfected wells. Blank wells of RPMI-only corrected for background in all samples. Samples were evaluated as per the manufacturer's instructions. Absorbance values from each well were measured at 490 nm on a microplate reader (POLARstar Omega, BMG LABTECH) and the % cytotoxicity was calculated using the following equation:

$$\frac{high\ control - low\ control}{sample - low\ control}\ x\ 100$$

In some experiments, cytotoxicity of macrophages induced by LPS (1 µg/mL) or LPS plus nigericin (20 µM) was also assessed.

**Western blots**
Differentiated THP-1 cells were seeded on 6-well plates at 1 x 106 cells/well. Isolates E104107R and E. cloacae ATCC13047 type-strain were selected for study. A HK control was included for isolate E104107R. The infection was carried out for 5 hours in total, 4 hrs infected with live isolates following nigericin (20 µM) for 1 hour as the positive control for pyroptosis. LPS-treated (1 ug/ml) THP-1 cells were used as inflammasome priming controls. The priming effect of LPS from the surface of Ecc isolates was also examined, including a control of HK infected and nigericin-treated cells was also included for both E104107R and ATCC13047. Conditions were established in triplicate. Cell-free supernatants from each triplicate well was transferred into the same pre-cooled microcentrifuge tube. THP-1 cells were washed three times with ice-cold PBS. 100 µl of Triton X-100 buffer (150 mM sodium chloride, 1 % Triton X-100, 50 mM Tris pH 8) was added to the first well of the triplicates, the lysate was collected and transferred to the next well and the process was repeated. The final 100 µl sample from each experimental condition was collected in the same pre-cooled microcentrifuge tube. Cell-free supernatants were concentrated prior to being loaded on the gel. This was achieved using StrataClean Resin. To quantify protein concentrations across samples and load equal amounts of protein on polyacrylamide gels, Bicinchoninic acid (BCA) protein assay was carried out. The lysate samples were sonicated for 10 minutes in a water bath sonicator to shear DNA and reduce sample viscosity, boiled at 95 °C for 10 minutes, centrifuged for 1 minute at 8000 rpm, resuspended in lysis buffer and loaded on the gel. The samples were electrophoresed using a Mini-Protean system (Bio-Rad). The proteins were transferred onto nitrocellulose membranes by wet transfer using a methanol-based buffer. The membranes were electrophoresed for 1 hour at 100 V. To confirm correct transfer of proteins, the samples were incubated with Revert 700 total protein stain as per the manufacturer's instructions. The membranes were washed once with TBST (20 mM Trizma base, 150 mM NaCl, pH 7.6 with 0.1% (v/v) Tween20) and blocked for 1 hour at room temperature with 5 % Blocker Casein in TBST. The membranes were washed with TBST prior to primary antibody incubation overnight at 4 °C with agitation. The membranes were washed with TBST and incubated at room temperature with the secondary goat anti- rabbit IRDye 800 cW (1:10000) (LI-COR) for 1 hour, in a dark chamber with agitation. The antibodies were made up in TBST. Finally, the membranes were washed again before being imaged on the LI-COR Odyssey scanner.

Table S1

| REAGENT | SOURCE | IDENTIFIER |
|---|---|---|
| **Antibodies** | | |
| Recombinant anti-cleaved N-terminal GSDMD; working dilution 1:1000 | Abcam | ab215203 |
| Monoclonal anti-Caspase-1; working dilution 1:1000 | Cell Signalling Technology | #3866D7F10 |
| Polyclonal anti-IL-1β; working dilution 1:1000 | Abcam | ab2105 |
| LAMP-1; working dilution 1:800 | Abcam | ab24170 |
| EEA1; working dilution 1:1142 | Invitrogen | PA1-063A |
| LC3B antibody kit for autophagy; working dilution 0.5 μg/mL | Invitrogen | L10382 |
| Rabbit IgG (Alexa488); working dilution 1:2000 | Invitrogen | A11034 |
| **Fluid Phase Marker** | | |
| Calcein Blue; working dilution 50 μg/mL | ThermoFisher Scientific | C1429 |
| Dextran Tetramethylrhodamine; working dilution 50 μg/mL | ThermoFisher Scientific | D1868 |
| Lysotracker; working dilution 250 μM | Invitrogen | L7526 |
| **Labelling in live bacteria** | | |
| 3-[[(7-Hydroxy-2-oxo-2*H*-1-benzopyran-3-yl)carbonyl]amino]-D-alanine hydrocholoride; HADA; working dilution 250 μM | Tocris Bioscience | 6647 |

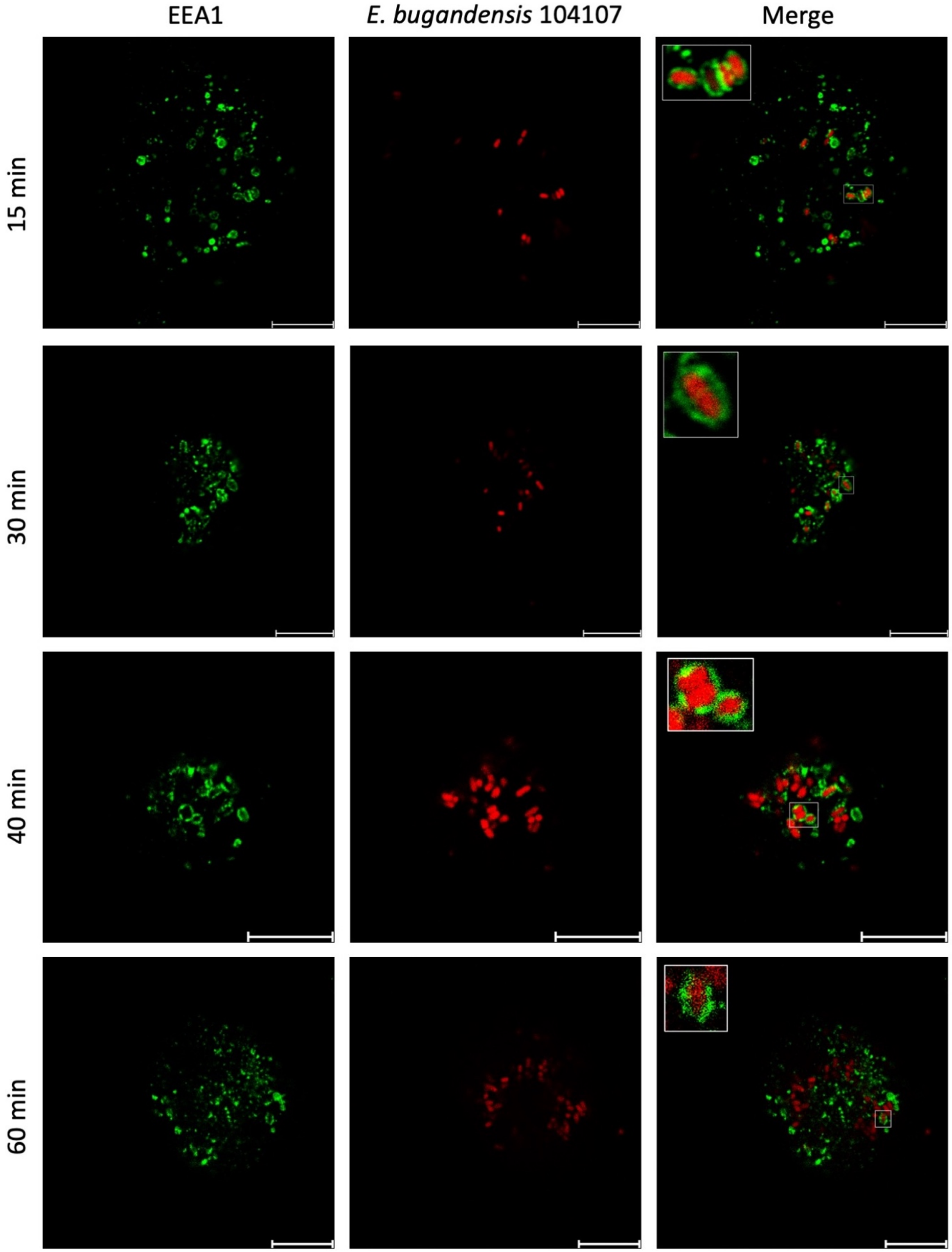


**Figure S1. Intracellular *E. bugandensis* 104107 bacteria traffic to EEA1-positive phagosomes in THP-1 cells.** Confocal microscopy of macrophages infected with *E. bugandensis* 104107 (pJT04). Bacteria appear in membrane vacuoles that acquire EEA1. Images taken with x 63 magnification on a Leica SP8 confocal microscope. MOI = 120. Scale bar = 10 µm. The quantitative results of these experiments are presented in Figure 1B.

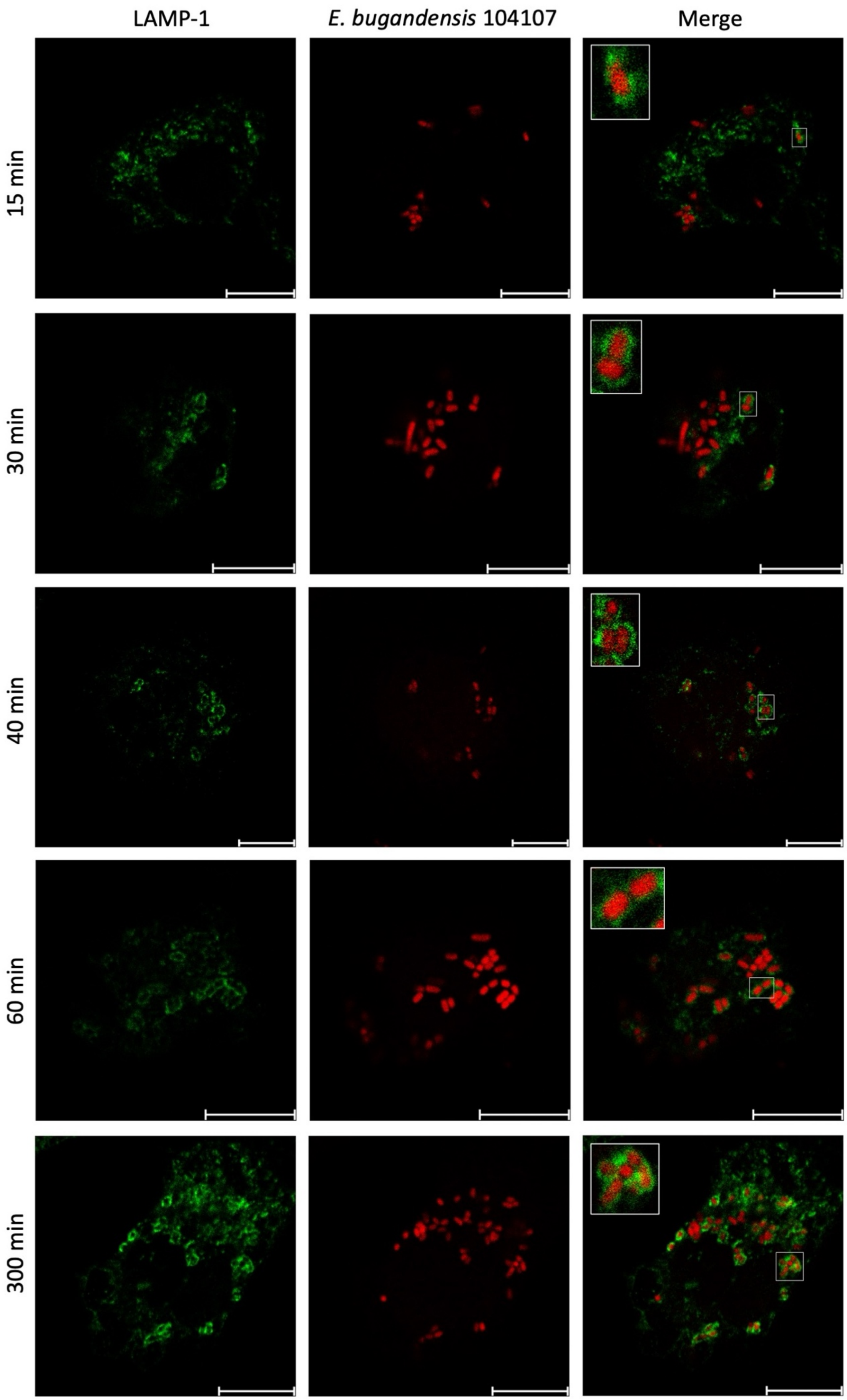
LAMP-1
E. bugandensis 104107
Merge
15 min
30 min
40 min
60 min
300 min

**Figure S2. Intracellular *E. bugandensis* 104107 bacteria traffic to LAMP1-positive phagosomes in THP-1 cells.** Confocal microscopy of macrophages infected with *E. bugandensis* 104107 (pJT04). Bacteria appear in membrane vacuoles that acquire LAMP1. Images taken with x 63 magnification on a Leica SP8 confocal microscope. MOI = 120. Scale bar = 10 μm. The quantitative results of these experiments are presented in Figure 1B.

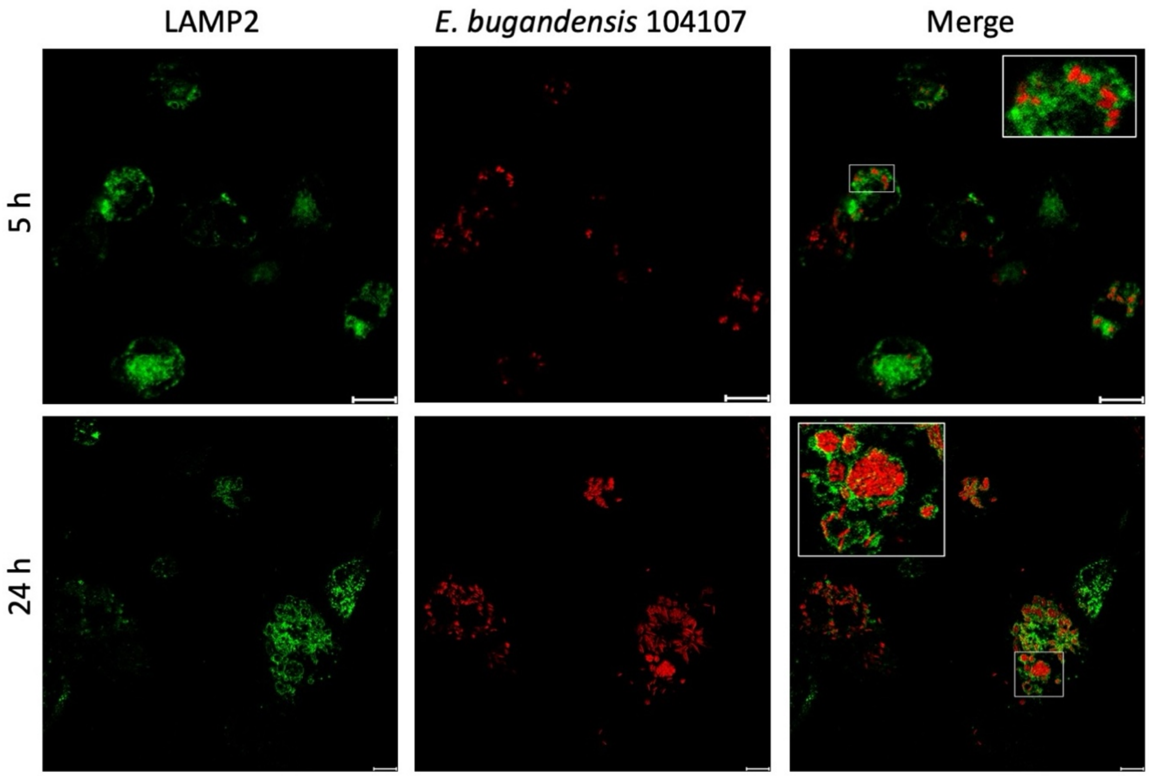


**Figure S3.** ***E. bugandensis*** **104107 bacteria traffic to LAMP2-rich compartments in THP-1 cells.** Transfected THP-1 macrophages expressing GFP-labelled LAMP2 were infected with live bacteria and examined at 5- and 24-hours post-infection. Images taken at x100 magnification on SP8 confocal microscope. MOI = 120. Scale bar = 10 µm. Insets indicate bacteria present in LAMP-2 decorated membrane vacuoles.

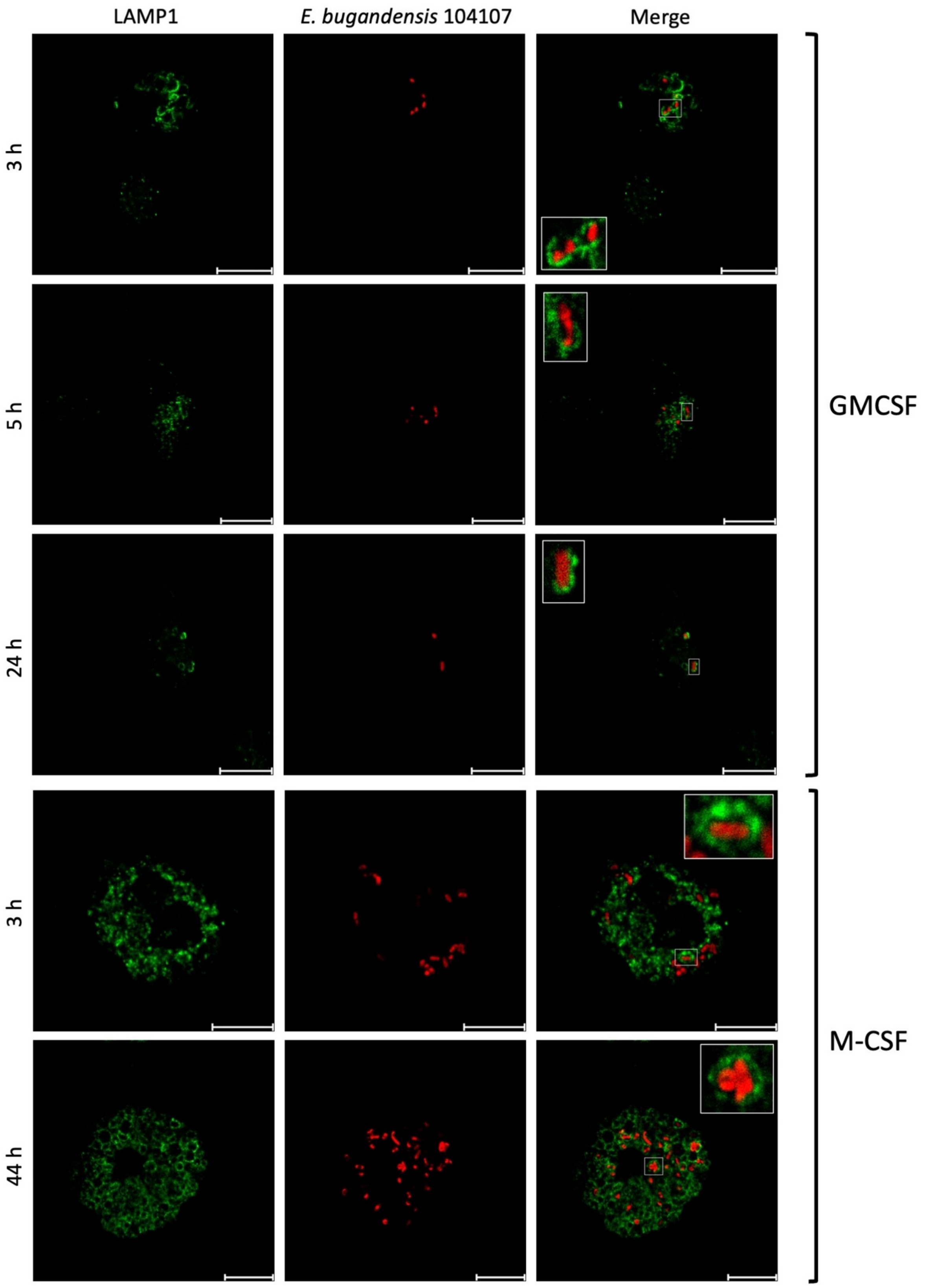


**Figure S4. *E. bugandensis* 104107 bacteria infecting HMDMs localise in LAMP1-rich compartments.** (top panel) GM-CSF differentiated HMDMs incubated with *E. bugandensis* 104107 for the indicated times colocalize with LAMP-1. (bottom panel) bacterial colocalisation in infected M-CSF differentiated HMDMs. All images were taken on a Leica SP8 confocal microscope with x 63. MOI = 40. Scale bar = 10 μm.

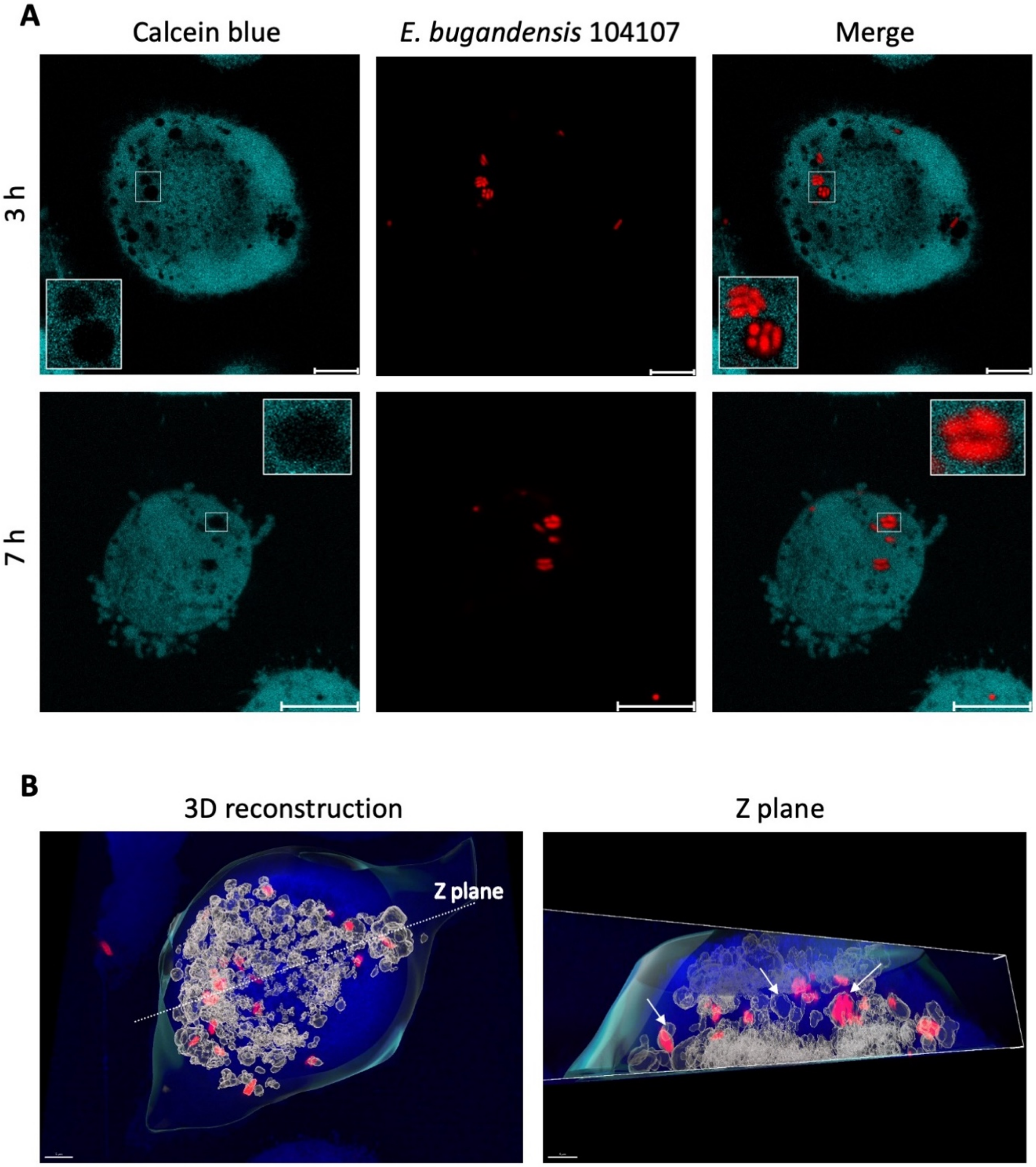


**Figure S5. Intracellular *E. bugandensis* 104107 remain within membrane vacuoles. A.** Live cell imaging of THP-1 macrophages infected with *E. bugandensis* 104107 (red fluorescence) at 3 hours and 7 hours post-infection. Cells were stained with calcein blue (original blue fluorescence pseudocoloured as cyan for better contrast), which does not enter membrane compartments. Images taken on a Leica Stellaris-5 confocal microscope with x 100 magnification. MOI = 15. **B.** 3D reconstruction model of an infected calcein blue-stained macrophage cell (left panel) indicating a Z plane optical section. The right panel indicates an optically inverted Z plane revealing that bacteria present in negative spaces corresponding to vacuoles. Other internal membrane cell compartments are detected without bacteria. Arrows indicate the location of the membranes.

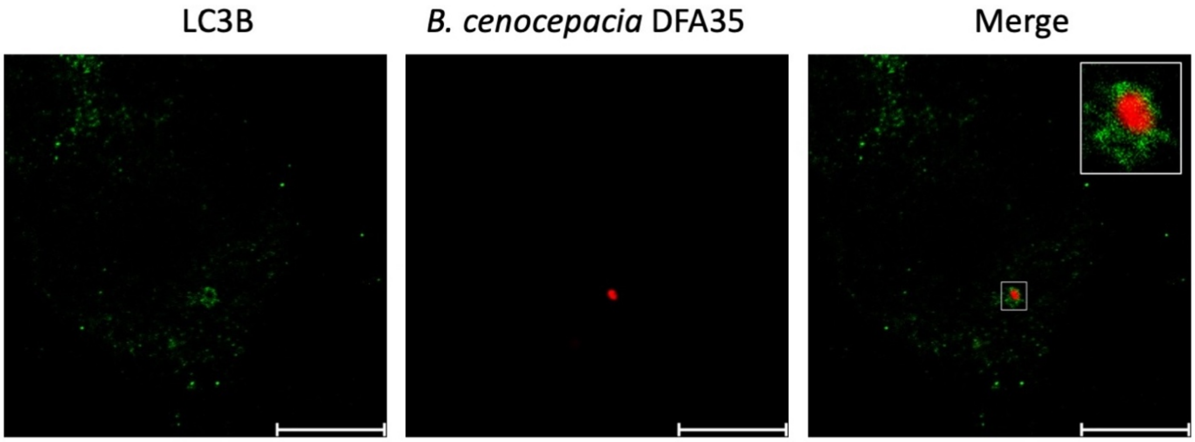


**Figure S6.** ***Burkholderia cenocepacia*** **colocalises with LC3B.** THP-1 macrophages were infected with *B. cenocepacia* strain DFA35 expressing mCherry and imaged at 3 hours post-infection. Images taken at x 100 magnification on SP8 confocal microscope. MOI = 30. Scale bar = 10 μm.

### Clinical isolates challenged with polymyxin B (PMB) reveal changes in their lipid A structures

These results complement and extend information provided in Fig. 5D of the manuscript. In addition to intracellualr survival in human macrophages, the *Enterobacter cloacae* complex isolates employed in this study were investigated for lipid A modifications in the absence or after challenge with 1 and 10 µg/mL PMB. The PMB challenge concentration depended on the intrinsinc PMB resistance of each isolate, as determined by broth microdilution in CA-MHB and published elsewhere [2,10]. Since resistance to polymyxins primarily depends on lipid A modifications [14], we inferred the lipid A structures of these isolates by comparing mass spectra profiles of isolated lipid A from pairs of untreated and PMB-treated isolates with using MALDI-TOF mass spectrometry (see details in the appendix, p. 3).

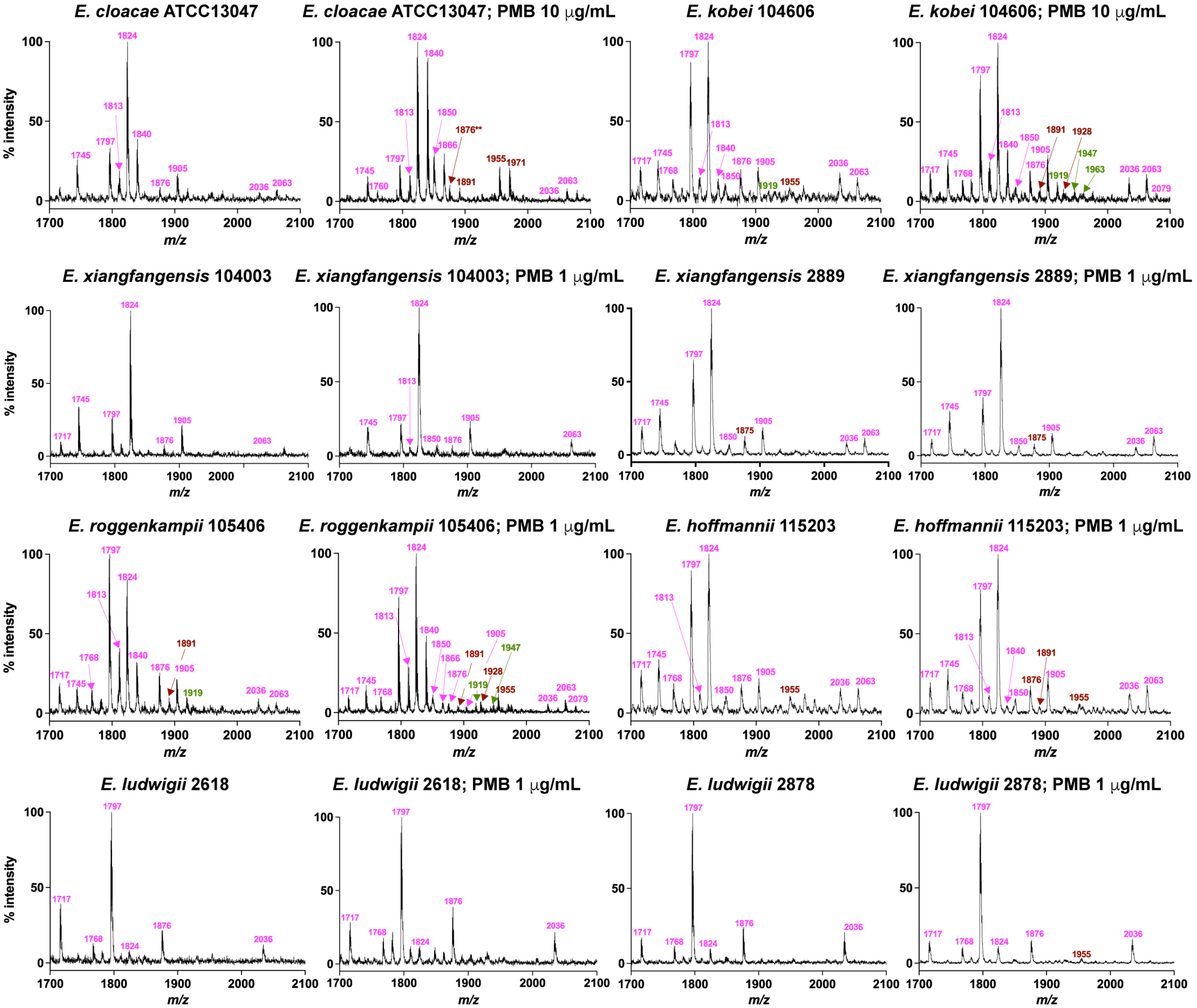


**Figure S7. Mass spectrometry profiles of lipid A isolated from clinical *Enterobacter* isolates.** The spectra in the absence of PmB revealed prominent peaks at *m/z* 1824 and 1797 (See Table S2 in appendix, p. 15) [2], corresponding to hexa-acylated lipid A forms, and 1387, which corresponds to a tetra-acyl form [2,10]. Similar peaks are found in *Klebsiella pneumoniae*, *Escherichia coli* and *Yersinia enterocolitica* [15]. Modified forms of these peaks were also present, such as the addition of a hydroxyl group (typically by the enzyme LpxO) (*m/z* 1840 to 1824, 1813 to 1797) and palmitate group (typically by PagP) (*m/z* 2063 to 1813) [16,17]. Peaks present in other isolates included *m/z* 1359 which is tetra-acyl like 1387 but with one C14 chain reduced in length to C12, *m/z* 1597 which corresponds to a palmitoylated or penta-acyl version of *m/z* 1359, and *m/z* 2036 which corresponds to a palmitoylated or hepta-acyl *m/z* 1797. Spectra obtained from PMB-challenged isolates displayed additional ion peaks which were either absent in the paired unchallenged isolate, or with significantly increased intensity in the presence of PmB (e.g., ion peak at *m/z* 1955, which coresponds to the addition of L-Ara4N [*m/z* +131 to the *m/z* 1824 hexa-acyl species]).

**Table S2. Lipid A composition deduced from MALDI-TOF mass spectrometry data.**

| Lipid A composition | Calculated *m/z* | Observed m/z |
|---|---|---|
| Hexa-acyl (4xC14:0(3-OH), 1x C14:0, 1x C12:0, 1P | 1716.17 | 1717 |
| Hexa-acyl (4xC14:0(3-OH), 2x C14:0, 1P | 1744.17 | 1745 |
| Hexa-acyl (4xC14:0(3-OH), 1x C14:0, 1x C14:0(3-OH), 1P | 1760.17 | 1760 |
| Hexa-acyl (4xC14:0(3-OH), 1x C14:0, 1x C12:0, 2P | 1797.14 | 1797 |
| Hexa-acyl (4xC14:0(3-OH), 1x C14:0, 1x C12:0(3-OH), 2P | 1813.14 | 1813 |
| Hexa-acyl (4xC14:0(3-OH), 2x C14:0, 2P | 1825.14 | 1824 |
| Hexa-acyl (4xC14:0(3-OH), 1x C14:0, 1x C14:0(3-OH), 2P | 1841.14 | 1840 |
| Hexa-acyl (4xC14:0(3-OH), 1x C14:0, 1x C16:0, 2P | 1852.14 | 1850 |
| Hexa-acyl (4xC14:0(3-OH), 1x C14:0(3-OH), 1x C16:0, 2P | 1868.14 | 1866 |
| Hexa-acyl (4xC14:0(3-OH), 2x C14:0, 1P, 1x L-Ara4N | 1875.17 | 1876 |
| Hexa-acyl (4xC14:0(3-OH), 1x C14:0, 1x C14:0(3-OH), 1P, L-Ara4N | 1892.17 | 1891 |
| Hexa-acyl (4xC14:0(3-OH), 2x C14:0, 3P | 1906.11 | 1905 |
| Hexa-acyl (4xC14:0(3-OH), 1x C14:0, 1x C12:0, 2P, 1x PEtN | 1920.14 | 1919 |
| Hexa-acyl (4xC14:0(3-OH), 1x C14:0, 1x C12:0, 2P, 1x L-Ara4N | 1928.14 | 1928 |
| Hexa-acyl (4xC14:0(3-OH), 2x C14:0, 2P, 1x PEtN | 1948.14 | 1947 |
| Hexa-acyl (4xC14:0(3-OH), 2x C14:0, 2P, 1x L-Ara4N | 1955.14 | 1955 |
| Hexa-acyl (4xC14:0(3-OH), 1x C14:0, 1x C14:0(3-OH), 2P, 1x PEtN | 1964.14 | 1963 |
| Hexa-acyl (4xC14:0(3-OH), 1x C14:0, 1x C14:0(3-OH), 2P, L-Ara4N | 1971.14 | 1971 |
| Hepta-acyl (4xC14:0(3-OH), 1x C14:0, 1x C12:0, 1x C16:0, 2P | 2036.34 | 2036 |
| Hexa-acyl (4xC14:0(3-OH), 2x C14:0, 1x C16:0, 2P | 2064.34 | 2063 |
| Hexa-acyl (4xC14:0(3-OH), 1x C14:0, 1x C14:0(3-OH), 1x C16:0, 2P | 2080.34 | 2079 |

C, carbon; P, phosphate; L-Ara4N, 4-amino-4-deoxy-L-arabinose; PEtN, phosphoethanolamine.

### The levels of virulence in *Galleria mellonella* do not correlate with PMB resistance

These results complement and extend information provided in Fig. 5D of the manuscript. In addition to intracellualr survival in human macrophages, the *Enterobacter cloacae* complex isolates employed in this study were invesgitated for their relative virulence in the *Galleria mellonella* moth larvae infection model (see methodology in appendix, p. 3). Data for *E. bugandensis* strains 104107 and 115227, and *E. ludwigii* 2618 were published elsewhere [10] and not included in the Figure S8. The data indicate that infected larvae displayed different levels of survival ranging from 30% (for *E. bugandensis* 104107[10] to 97%. The Kaplan-Meier survival curves were analysed by log rank by multiple comparisons among isolates and to compare against the mock infected control (PBS). Two groups of isolates were clearly distinguishable: those displaying probability of survival between 73-97% (low virulence) and those with survival between 30-40%. The latter included *E. bugandensis* 104107[10] and *E. roggenkampii* 105406 (Figure S8). From these results, we concluded that the relative virulence of each isolate was strain dependent and could not be directly correlated with the level of PMB susceptibility, lipid A modifications, or the ability of the isolates to survive within human macrophages. This suggests that other factors may play a role in determining the infectivity of the isolates in the *G. mellonella* model.

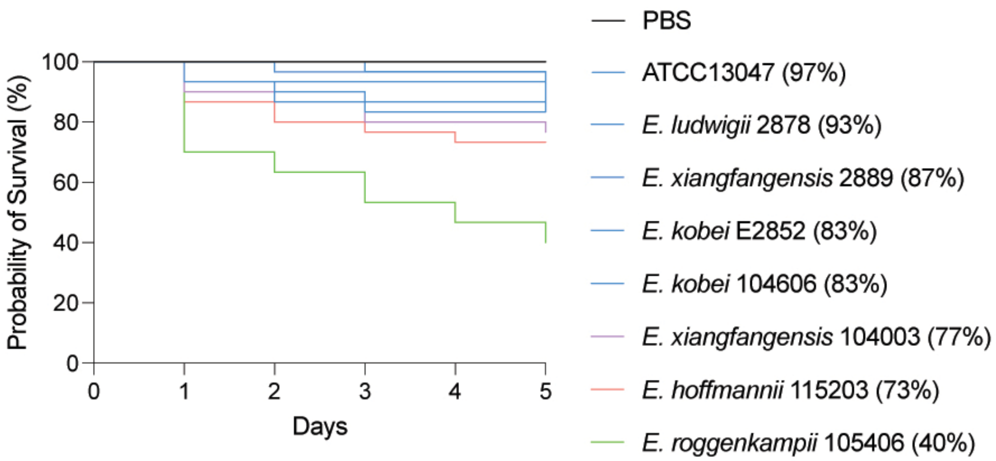


**Figure S8. The levels of relative virulence of *Enterobacter* clinical isolates in the *Galleria mellonella* infection model.** The Kaplan-Meier survival curves were analysed by log rank by multiple comparisons among isolates and to compare against the mock infected control.

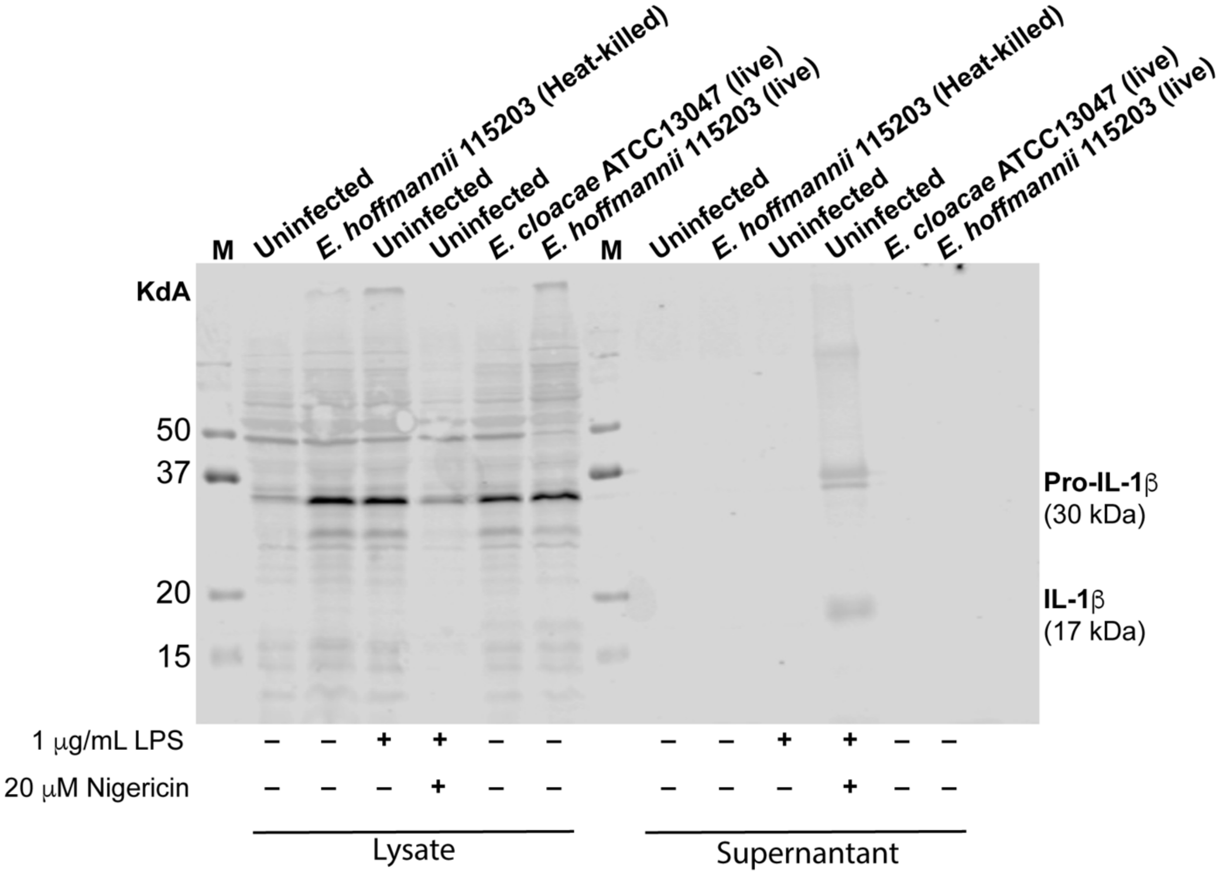


**Figure S9. THP-1 macrophages infected with *E. hoffmannii* 115203 and *E. cloacae* ATCC13047 do not induce cleavage of Pro-IL-1β.** THP-1 macrophages were infected with the bacteria and incubated for 5 hours. Uninfected macrophages were also examined as a control. In some cases, uninfected macrophages were treated with Heat-killed *E. hoffmannii* 115203, LPS (priming signal) or LPS + Nigericin (positive control to induce pyroptosis and cleavage of Pro-IL-1β). The cells lysates from macrophages not exposed to bacteria show smaller amounts of Pro-IL-1β, while lysates of macrophages exposed to bacteria, LPS or LPS + Nigericin show more Pro-IL-1β expression. IL-1β was only found in the supernatant of macrophages treated with LPS + Nigericin, suggesting that either heat-killed or live *Enterobacter* bacteria prime macrophages but do not induce pyroptosis.
